# The pathogenic E139D mutation stabilizes a non-canonical active state of the multi-domain phosphatase SHP2

**DOI:** 10.1101/2025.07.02.662799

**Authors:** Anne E. van Vlimmeren, Ziyuan Jiang, Deepti Karandur, Anya T. Applebaum Licht, Neel H. Shah

## Abstract

Dysregulation of the phosphatase SHP2 is implicated in various diseases, including congenital disorders and cancer. SHP2 contains two phosphotyrosine-recognition domains (N-SH2 and C-SH2) and a protein tyrosine phosphatase (PTP) domain. The N-SH2 domain is critical for SHP2 regulation. In the auto-inhibited state, it binds to the PTP domain and blocks the active site, but phosphoprotein engagement destabilizes the N-SH2/PTP domain interaction, thereby exposing the active site. Many disease mutations in SHP2 are at the N-SH2/PTP interface, and they hyperactivate SHP2 by disrupting auto-inhibitory interactions. The activating E139D mutation represents an exception to this mechanism, as it resides in the C-SH2 domain and makes minimal interactions in auto-inhibited and active state crystal structures. In this study, using AlphaFold2 modeling and molecular dynamics simulations, we identify an alternative active conformation of SHP2, in which Glu139 interacts with Arg4 and Arg5 on the N-SH2 domain to stabilize a novel N-SH2/C-SH2 interface. Using double mutant cycles, we show that this active state is further stabilized by the E139D mutation. Finally, we demonstrate that the E139D mutation enforces an active conformation with distinct phosphoprotein binding preferences from canonical hyperactive mutants. Thus, our study reveals a novel mechanism for SHP2 dysregulation.

## Introduction

SHP2 is a protein tyrosine phosphatase with diverse and critical roles in cell signaling and human physiology (Scheiter *et al*, 2024; Lauriol *et al*, 2015; Salmond & Alexander, 2006). The importance of SHP2 function is further underscored by the many disease-associated missense mutations that have been identified in *PTPN11*, the gene encoding SHP2. The resulting amino acid substitutions are primarily known for dysregulating SHP2 to cause the developmental disorders Noonan Syndrome and Noonan Syndrome with Multiple Lentigines (Edouard *et al*, 2010; Tartaglia *et al*, 2006). However, SHP2 mutations can also drive cancer: 40% of juvenile myelomonocytic leukemia (JMML) patients have SHP2 mutations (Gupta *et al*, 2021), and less frequent mutations are also found in acute myeloid leukemia (AML), acute lymphoid leukemia (ALL), and solid cancers (Fobare *et al*, 2022; Estey, 2018; Bentires-Alj *et al*, 2004).

The functional consequences of most disease-associated mutations in SHP2 can be readily understood by examining its structure. SHP2 has three globular domains: a catalytic protein tyrosine phosphatase (PTP) domain, and two SH2-domains that regulate SHP2 activation and localization (Hof *et al*, 1998). The N-SH2 and PTP domains make extensive interactions that keep SHP2 in an auto-inhibited conformation in the apo state (**Figure 1A**). Engagement of the N-SH2 domain by phospho-ligands disrupts auto-inhibitory interactions between these two domains, which induces a conformational change that makes the catalytic site accessible to substrates (Hof *et al*, 1998). Many mutations in SHP2-related disorders are found at this auto-inhibitory interface, and they hyperactivate SHP2 by disrupting auto-inhibition. Several other mutations are located outside of this interface. Some indirectly disrupt auto-inhibition by destabilizing the N-SH2 core or by altering the positioning of loops that can sterically block the N-SH2/PTP interaction (Jiang *et al*, 2025). Others dysregulate SHP2 by altering substrate or activator recognition (Martinelli *et al*, 2008; Toto *et al*, 2020; Zhang *et al*, 2020; van Vlimmeren *et al*, 2024). Several mutations outside of the N-SH2/PTP interface activate SHP2 through currently ambiguous mechanisms.

**Figure 1.**
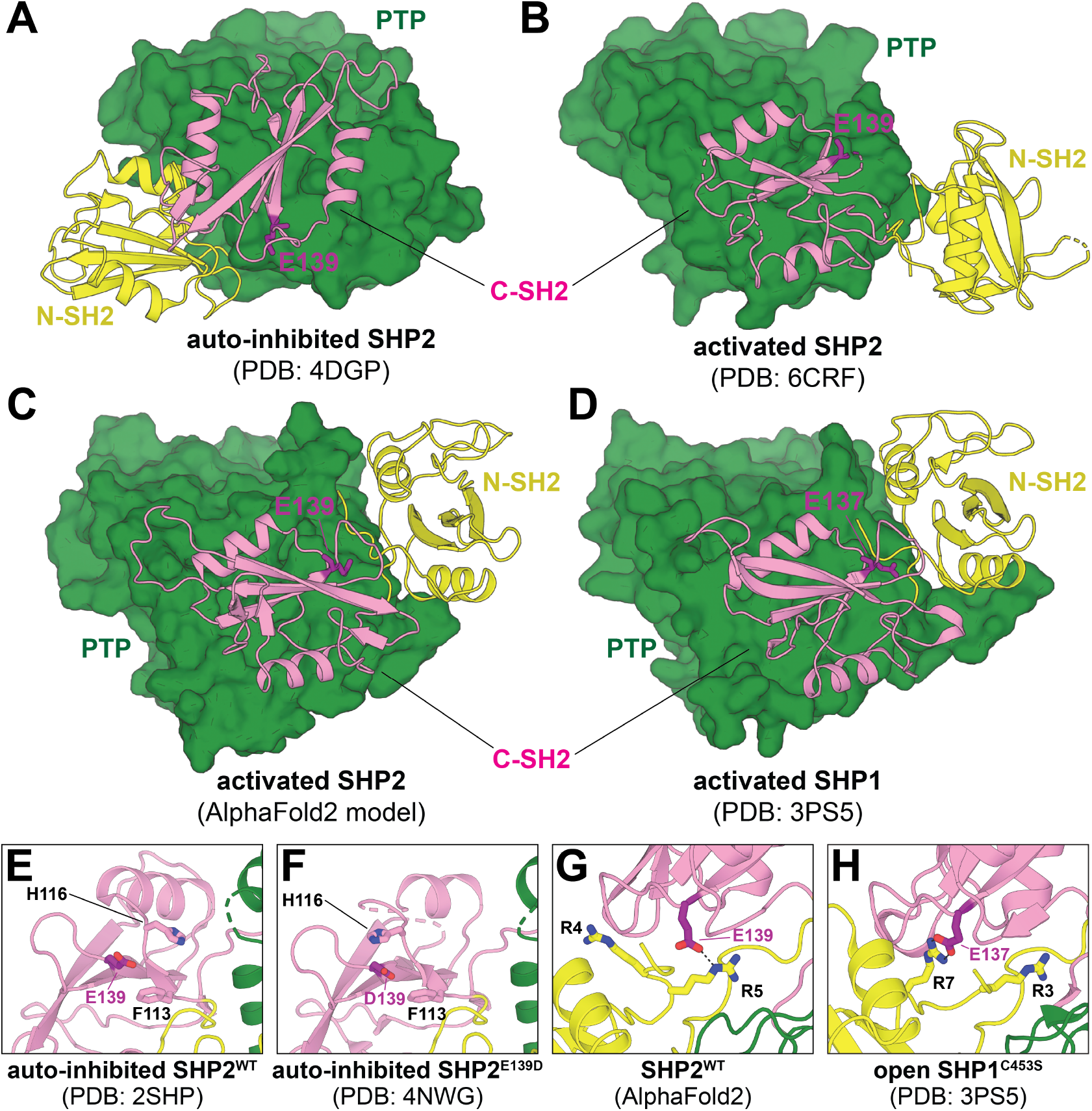
Auto-inhibited and open conformations of SHP2. (**A**) Crystal structure of SHP2^WT^ in an auto-inhibited conformation, maintained by auto-inhibitory interactions between the N-SH2 (*yellow*) and PTP domain (*green*) (PDB code 4DGP). (**B**) Crystal structure of SHP2^E76K^ in an open conformation. SH2 domains are rotated and N-SH2 domain is in a dramatically different position relative to the PTP domain (PDB code 6CRF). (**C**) AlphaFold2 model of SHP2 in an open conformation in which the N-SH2 domain adopts a distinct position and orientation relative to PDB structure 6CRF. (**D**) Crystal structure of SHP1^C453S^ in an open conformation that largely resembles the AlphaFold2 SHP2 state (PDB code 3PS5). (**E**) Glu139 in the auto-inhibited state does not make any interdomain contacts (PDB code 2SHP). (**F**) Asp139 in auto-inhibited state does not make any interdomain contacts (PDB code 4NWG). (**G**) Glu139 interacts with Arg5 in the AlphaFold2 model of SHP2. (**H**) Glu137 interacts with Arg7 in the open state structure of SHP1 (PDB code 3PS5).

Much of our understanding of the catalytically-active conformation of SHP2 comes from a single crystal structure of the JMML mutant SHP2^E76K^ (**Figure 1B**, PDB code 6CRF). This mutation severely impedes auto-inhibitory interactions between the N-SH2 and PTP domains (Hof *et al*, 1998; LaRochelle *et al*, 2018). In the published crystal structure, SHP2^E76K^ inhabits an “open conformation”, where the C-SH2 domain pivots on the face of the PTP domain, and the N-SH2 domain is displaced by more than 120° around this C-SH2 pivot axis, leaving the catalytic site exposed and accessible to substrates. In a recent report from our group (Jiang *et al*, 2025), we described an AlphaFold2 (AF2, (Jumper *et al*, 2021)) model of wild-type SHP2 in which the C-SH2 domain makes a similar pivot relative to the PTP domain, but the N-SH2 domain is further behind the PTP domain and rotated relative to the C-SH2 domain (**Figure 1C**). Some key features of the C-SH2/PTP interface are conserved between this AF2 model and the 6CRF crystallographic active state, such as a stabilizing ion pair between R111 and E249 (Kim *et al*, 2024), and both structures are compatible with the observed mutational sensitivity at the C-SH2/PTP interface seen in our recent deep mutational scanning study (Jiang *et al*, 2025). Notably, the SHP2 AF2 model resembles a crystal structure of the closely-related phosphatase SHP1 in an active conformation (PDB code 3PS5), which has similar positioning and orientation of the SH2 domains relative to the PTP domain and to one another (**Figure 1D**) (Wang *et al*, 2011). Recent studies have suggested that the conformation observed in the SHP1 open structure, and not the conformation in the SHP2^E76K^ crystal structure, is the state that is adopted when SHP2 is bound to certain bis-phosphorylated activators (Marasco *et al*, 2020; Perla *et al*, 2025). Additionally, there is growing evidence from small angle X-ray scattering, single molecule fluorescence, and computation that SHP2 can exist in more than one active state (Hayashi *et al*, 2017; Pádua *et al*, 2018; Calligari *et al*, 2021; Tao *et al*, 2021; Anselmi & Hub, 2023). Refined structural models of the various states in this ensemble may shed light on mechanistic details of SHP2 activation and mutational effects that cannot currently be explained by existing crystal structures.

SHP2^E139D^ is a pathogenic mutant that exists in both Noonan Syndrome and JMML and has been shown to have increased basal catalytic activity (Keilhack *et al*, 2005; Tartaglia *et al*, 2006; Martinelli *et al*, 2008; van Vlimmeren *et al*, 2024). The E139D mutation has also been shown to enhance the activation of SHP2 by phosphopeptides (Keilhack *et al*, 2005; Tartaglia *et al*, 2006; Martinelli *et al*, 2008). Interestingly, Glu139 is not located at the auto-inhibitory interface but lies in the C-SH2 domain, and its mechanism of dysregulation remains unknown. Based on the previously determined auto-inhibited structures of SHP2^WT^, there is no obvious role for Glu139 (**Figure 1A,E**) (Yu *et al*, 2013), and the side chain of this residue was not resolved in the open state structure of SHP2^E76K^ (**Figure 1B**) (LaRochelle *et al*, 2018). Moreover, in a crystal structure of SHP2^E139D^, which is in the auto-inhibited state, no acquired role was found for Asp139, although there was some minor local side chain repositioning (**Figure 1F**) (Qiu *et al*, 2014). Interestingly, in the open state AlphaFold2 model of SHP2, Glu139 interacts with Arg5 in at the N-terminus of the N-SH2 domain (**Figure 1G**). Furthermore, the analogous residue in SHP1, Glu137, engages with Arg7 on the N-SH2 domain in the active state SHP1 structure (**Figure 1H**) (Wang *et al*, 2011). These observations raise the possibility that the E139D mutation exerts its effects not by disrupting auto-inhibition, but instead by stabilizing an active state of SHP2. This is supported by the finding that E139D is the only substitution at this residue that is hyperactivating in our previously reported deep mutational scan of SHP2 (**Supplementary Figure 1**) (Jiang *et al*, 2025).

Here, we describe the biochemical characterization of key residues in an interaction network involving Glu/Asp139, identified via molecular dynamics (MD) simulations of the AF2-predicted open state. We demonstrate that the hyperactivation of SHP2^E139D^ is likely mediated by stabilizing interactions between Asp139, Arg5, and potentially Arg4 in this non-crystallographic open state. We also provide evidence that SHP2^E139D^ has altered phosphoprotein binding preferences compared to other SHP2 variants, potentially due to geometric constraints imposed by the stabilization of a specific active state conformation. Our results shed light on a potential molecular mechanism underlying the pathogenicity of the E139D mutation and further support the conformational ensemble model for active SHP2.

## Results

### Glu/Asp139 interacts with N-terminal Arg residues in an alternative SHP2 open conformation

To compare interdomain orientation and dynamics in different open conformations of SHP2, we revisited our recently published triplicate 2.5 µs MD simulations starting from the open conformation crystal structure (PDB code 6CRF) and the AF2 model, both with the SHP2^WT^ sequence (Jiang *et al*, 2025). On the timescale of these simulations, both SHP2^WT^/6CRF and SHP2^WT^/AF2 sample a range of positions and orientations of the N-SH2 domain relative to the C-SH2 and PTP domains, but neither open state ever fully transitions to the other open state (**Figure 2A**). We quantified this movement by measuring an angle defined by N-SH2 position in the auto-inhibited state (PDB code 4DGP), a pivot point on the C-SH2 domain, and N-SH2 position in the open state simulations (**Supplementary Figure 2A**). In general, the N-SH2 domain retained a larger rotation angle in the SHP2^WT^/AF2 simulations relative to the SHP2^WT^/6CRF simulations (**Figure 2B**). The C-SH2 domain itself also rotates between the auto-inhibited and open states, while sitting on the face of the PTP domain. We quantified this rotation by measuring an angle defined by two vectors on the PTP and C-SH2 domains that are nearly parallel in the auto-inhibited state and substantially unaligned in the active states (**Supplementary Figure 2B**). On average, the C-SH2 domain rotation was slightly larger in SHP2^WT^/AF2 simulations than in the SHP2^WT^/6CRF simulations (**Figure 2C**).

**Figure 2.**
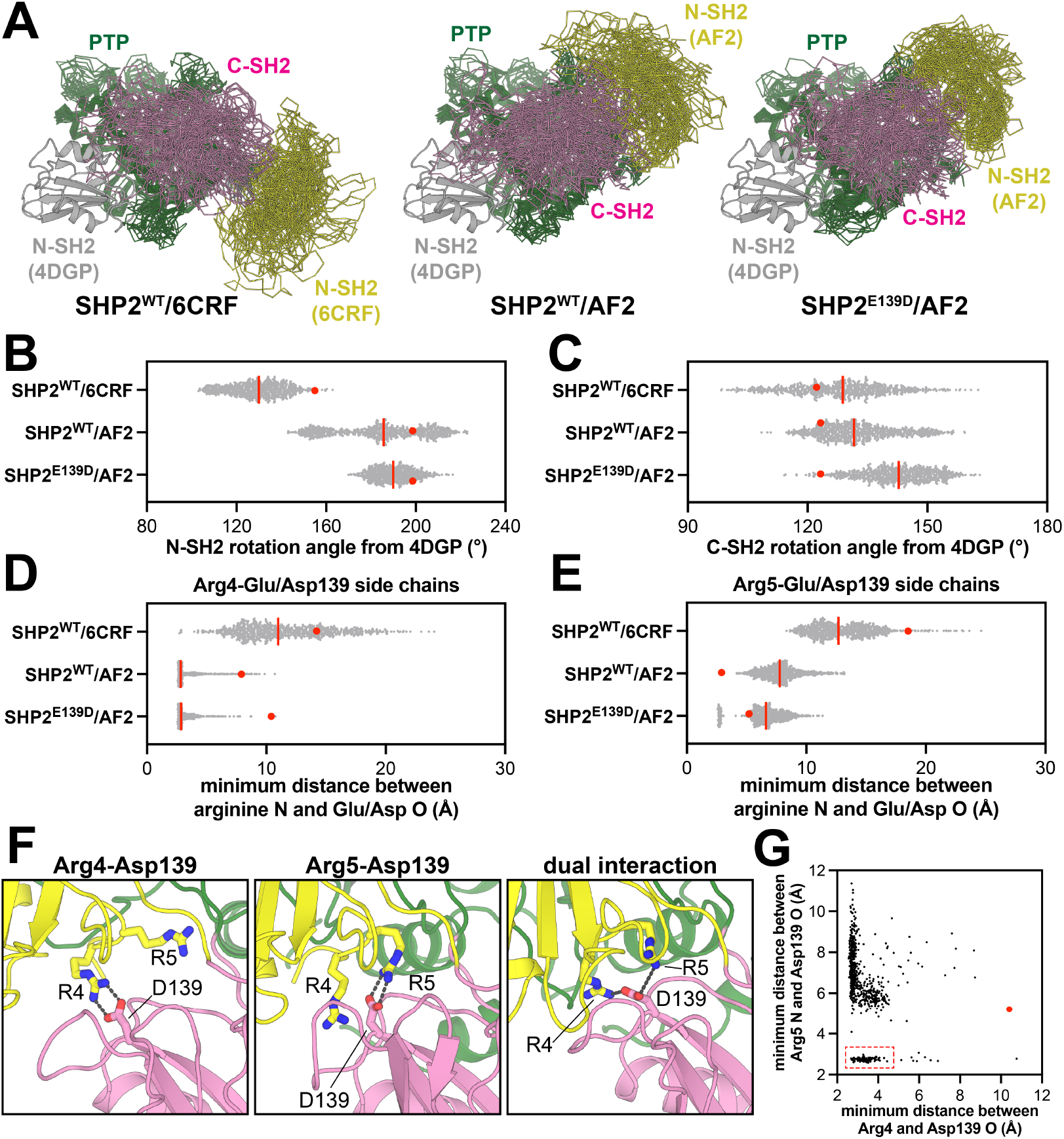
A non-crystallographic open conformation of SHP2 is stabilized by Glu/Asp139 electrostatic interactions in molecular dynamics simulations. (**A**) Representative sampling of structures from MD simulation trajectories of SHP2^WT^/6CRF (*left*), SHP2^WT^/AF2 (*middle*), and SHP2^E139D^/AF2 (*right*). Yellow, pink, and green represent N-SH2, C-SH2, and PTP domains, respectively. N-SH2 domain positioning in the closed conformation (PDB code 4DGP) is shown in gray as a reference, and all states are aligned over just the PTP domain residues. (**B**) Distribution of N-SH2 rotation angles from the closed conformation (PDB code 4DGP) to the positions seen in the SHP2^WT^/6CRF, SHP2^WT^/AF2, and SHP2^E139D^/AF2 simulations. (**C**) Same as (**B**) but measuring C-SH2 rotation. (**D**) Minimum distance between Arg4 side-chain N atoms and Glu/Asp139 side-chain O atoms in the SHP2^WT^/6CRF, SHP2^WT^/AF2, and SHP2^E139D^/AF2 simulations. (**E**) Same as (**D**), but for Arg5. In panels (**B**) to (**E**), the red line indicates the median value of the distribution, and the red dot denotes the measurement in the starting model used for that simulation. (**F**) Renderings of structures from SHP2^E139D^/AF2 simulations, highlighting Arg4 and Arg5 interactions with Asp139. (**G**) Concurrent Arg4-Asp139 and Arg5-Asp139 distances across SHP2^E139D^/AF2 simulations, showing a population of frames where both interactions occur simultaneously (dashed red box). The red dot denotes the measurement in the starting model used for that simulation.

As a consequence of these different interdomain orientations, Glu139 was substantially more buried in SHP2^WT^/AF2 simulations when compared with SHP2^WT^/6CRF simulations (**Supplementary Figure 3**). Another consequence of this change in interdomain orientations was that Arg4 and Arg5 at the N-terminus were proximal to Glu139 in the SHP2^WT^/AF2 simulations. Surprisingly, although Arg5 and Glu139 form an ion pair in the AlphaFold2 model used as a starting structure (**Figure 1G**), local rearrangements during the simulations led to Arg4 forming a more persistent ion pair with Glu139, with Arg5-Glu139 interactions happening less frequently (**Figure 2D,E**). This occurs in all three, independent trajectories, suggesting that the AlphaFold2 model is a metastable state that relaxes over the course of the simulations. By contrast, the two arginine residues rarely came in close proximity to Glu139 in the SHP2^WT^/6CRF simulations (**Figure 2D,E**). The Glu139-Arg4 and Glu139-Arg5 interactions are similar to the interaction between Glu137 and Arg7 in the open conformation crystal structure of SHP1 (**Figure 1H**) (Wang *et al*, 2011). We hypothesize that these electrostatic interactions play a role in stabilizing the SHP1 and AF2 active conformations and are not relevant in the distinct 6CRF active conformation.

To evaluate how the E139D mutation might alter interactions at the N-SH2/C-SH2 interface, we conducted triplicate 2.5 µs MD simulations starting from our original AF2 active state model, but with the Glu139 mutated to Asp. Asp139 remained buried at the N-SH2/C-SH2 interface in these simulations (**Supplementary Figure 3**). However, when compared to SHP2^WT^/AF2, the SHP2^E139D^/AF2 simulations showed decreased N-SH2 dynamics and an increase in the C-SH2 rotation angle (**Figure 2A-C**). We hypothesize that the shorter side chain of Asp139 stabilizes the further rotated SH2 domains. At the N-SH2/C-SH2 interface, Asp139 formed ion pairs with both Arg4 and Arg5, as also seen in the SHP2^WT^/AF2 simulations, however the Asp139-Arg5 interaction frequency increased in the mutant simulations when compared to Glu139-Arg5 in the wild-type simulations (**Figure 2D-F**). A small fraction of the time, Asp139 simultaneously engaged Arg4 and Arg5, suggesting that these interactions are not necessarily mutually exclusive (**Figure 2F,G**). Given the short timescale of the simulations, the relative contributions of Arg4 and Arg5 cannot be accurately determined. Regardless, these simulations suggest that Glu139 makes interdomain electrostatic interactions in the AF2-predicted SHP2 active conformation that are further enhanced with the E139D mutation, and the stabilization of this state may explain the activating effect seen for this mutation.

### The activating role of Glu/Asp139 is dependent on Arg4 and Arg5

To evaluate the functional effects of Arg4/Arg5 salt bridges with Glu/Asp139, we experimentally characterized single mutations at these residues, as well as a series of double and triple mutants (**Supplementary Table 1**). We first compared the catalytic efficiencies (*k*_cat_/*K*_M_ values) of SHP2^WT^, SHP2^E139D^, SHP2^R4A+R5A^, and SHP2^R4A+R5A+E139D^ using 6,8-difluoro-4-methylumbelliferyl phosphate (DiFMUP), a commonly used fluorogenic phosphatase substrate. As expected, the E139D mutation increased SHP2 basal activity 7-fold. However, in the background of the R4A+R5A double mutation, E139D no longer altered SHP2 activity (**Figure 3A**). These results demonstrate that the activating effect of Asp139 is dependent on the presence of Arg4 and/or Arg5.

**Figure 3.**
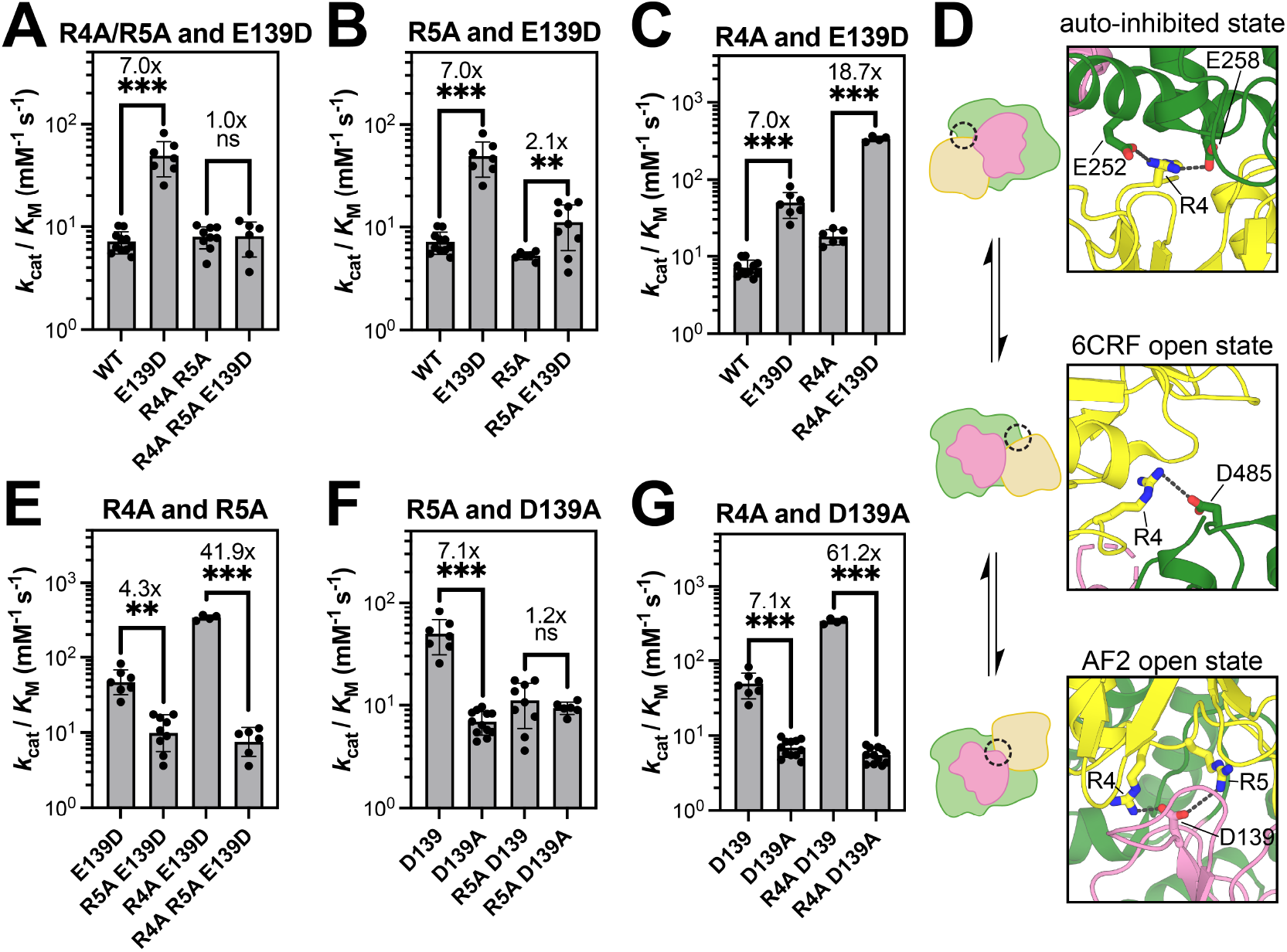
Coupling between Arg4, Arg5 and Glu/Asp139. (**A**) The activating effect of the mutation E139D is dependent on the presence of Arg4/Arg5. (**B**) The R5A mutation decreases the activating effect of E139D. (**C**) The R4A mutation enhances the activating effect of E139D. (**D**) Arg4 participates in distinct electrostatic interactions in the auto-inhibited state (PDB code 4DGP), the 6CRF open state (PBD code 6CRF), and the AF2 open state (representative frame from E139D MD trajectory. (**E**) The inactivating effect of the R5A mutation in an E139D background is significantly enhanced by the R4A mutation. (**F**) The inactivating effect of the D139A mutation is lost in an R5A background. (**G**) The inactivating effect of the D139A mutation is amplified in an R4A background. In all panels, statistical significance was assessed by Welch’s two-tailed t test. ns = not significant, * denotes p < 0.05, ** denotes p < 0.01, *** denotes p < 0.001.

To dissect the roles of Arg4 and Arg5, we performed additional mutant cycles analyzing R4A and R5A separately with E139D. The R5A mutation on its own had minimal effect on SHP2 activity, but it significantly weakened the activating effect of E139D from 7-fold to 2.1-fold (**Figure 3B and Supplementary Figure 4A**). This is consistent with Arg5 playing an important role in stabilizing the active AF2 state via Asp139. Interestingly, we found that the R4A mutation mildly increased SHP2 activity on its own and actually amplified the activating effect of E139D from a 7-fold to 18.7-fold enhancement in activity (**Figure 3C and Supplementary Figure 4B**). This result appears to contradict the significant role that the Arg4-Asp139 interaction plays in the AF2-state MD simulations – removing the Arg4 side chain should diminish the E139D effect, as seen for Arg5 (**Figure 3B**). This anomalous effect of the R4A mutation could be explained by the fact that Arg4 actually stabilizes multiple states of SHP2. In the auto-inhibited state, it interacts with Glu258 and potentially Glu252 (Hof *et al*, 1998; Yu *et al*, 2013), while in the 6CRF open state, it interacts with Glu485 (LaRochelle *et al*, 2018) (**Figure 3D**). Thus, while the R4A mutation would ablate the Arg4-Asp139 interaction seen in our AF2 state simulations, it would also destabilize competitive conformational states. Depending on the relative importance of Arg4 in each of these conformational states, it is possible that the R4A mutation might enhance occupancy of the AF2 state, which would now be more dependent on the Arg5-Asp139 interaction. Indeed, in an E139D background, the R5A mutation only causes a 4.3-fold reduction in activity, but in an R4A+E139D background, the R5A mutation causes a 41.9-fold reduction in activity (**Figure 3E**). From these experiments, it is clear that Arg5 plays a key role in mediating the activating effect of E139D, corroborating our AF2 model and simulations. However, the precise function of Arg4 cannot be unambiguously determined from activity measurements.

Our AF2 model and MD simulations not only suggest how the E139D mutation can stabilize an active SHP2 conformation, but they also suggest that Glu139 might be able to engage Arg4 and Arg5 in SHP2^WT^, albeit to a lesser extent than Asp139 in SHP2^E139D^. To investigate the interaction between Arg4, Arg5, and Glu139 in SHP2^WT^, we further performed mutant cycle analyses measuring activity levels of SHP2^R4A^, SHP2^R5A^, SHP2^E139A^, and the corresponding double mutants. Consistent with our deep mutational scanning data (**Supplementary Figure 1**) (Jiang *et al*, 2025), the E139A mutation did not have a measurable impact on catalytic activity on its own (**Supplementary Figure 5A**), indicating that the ability of Glu139 to stabilize an active state in SHP2^WT^ is likely negligible. When the E139A mutation was made in an R4A background, however, we observed a 3.2-fold reduction in activity (**Supplementary Figure 5A**). As noted above, outside of its potential role in the AF2 state, Arg4 also stabilizes alternate conformations, and so removal of Arg4 may enhance the population of SHP2 in the AF2 state. Thus, in this context, an effect of the E139A mutation can be detected. On the other hand, the R5A mutation caused a very modest 1.8-fold enhancement in the effect of the E139A mutation, suggesting that Glu139 and Arg5 are not energetically coupled in SHP2^WT^ (**Supplementary Figure 5B**). These data indicate that for SHP2^WT^, the active AF2 conformation is not significantly stabilized by interactions between Glu139 and Arg4/Arg5.

Finally, we compared constructs with E139D to those with E139A to analyze the impact of removing an Asp side chain from SHP2^E139D^. The “D139A” mutation (i.e. comparison of E139D to E139A) inactivated SHP2 7.1-fold, and this negative effect was fully lost in an R5A background, substantiating the importance of the Asp139-Arg5 ion pair (**Figure 3F and Supplementary Figure 5C**). By contrast, the 7.1-fold reduction in activity with the D139A mutation became a 61.2-fold reduction in activity in an R4A background (**Figure 3G and Supplementary Figure 5D**). Taken together, our data clearly demonstrate that there is an interplay between Arg4, Arg5, and Glu/Asp139 that governs the stability of the active AF2 state. Our MD simulations and mutant cycle analysis unambiguously show that Arg5 mediates the activating effect of the E139D mutation. However, the role of Arg4 is somewhat complicated. Our MD simulations point to a significant role for this residue in stabilizing the AF2 open conformation, however its role in other conformational states convolutes the mutant cycle analysis (**Figure 3D**). The fact that the R4A mutation sensitizes SHP2 to mutation at Arg5 suggests that these two arginine residues cooperate to stabilize the AF2 active state (**Figure 3E**). When Arg4 is removed, this state becomes highly dependent on Arg5. Finally, we note that there may be alternative explanations for our mutant cycle data that could be revealed through further structural analyses on Arg4 or Arg5 mutants.

### The activating E76K and E139D mutations have distinct effects on SHP2 structure

The E76K mutation serves as the canonical example of an activating SHP2 mutation at the N-SH2/PTP interface. SHP2^E76K^ is one of the most active SHP2 mutants, it can enhance basal activity against DiFMUP over 100-fold, and it cannot be further activated by phospho-peptide ligands, suggesting that it does not appreciably occupy the auto-inhibited state (LaRochelle *et al*, 2018; Jiang *et al*, 2025). Disruption of the auto-inhibited conformation by this mutation has been extensively characterized, and indeed, SHP2^E76K^ provided the first crystal structure of an open SHP2 conformation (PDB code 6CRF) (LaRochelle *et al*, 2018). Importantly, although SHP2^E76K^ has been crystallized in an open conformation, there is little direct evidence that this mutation significantly stabilizes that particular crystallographic state or any other open conformation of SHP2, as opposed to merely destabilizing the auto-inhibited state. One computational study suggested that, in SHP2^E76K^, Lys76 can make modest interactions with the PTP domain in the 6CRF state, but these have not been experimentally validated (Liu *et al*, 2023). By contrast, accumulating evidence suggests that the active state crystal structure of SHP2^E76K^ represents just one of multiple open conformations that SHP2 can access. Early crystallization attempts with full-length SHP2^E76K^ failed, attributed to the flexibility of the N-SH2 domain (Pádua *et al*, 2018). Subsequent experimental work confirmed alternative N-SH2 positions, supporting the existence of open conformations distinct from that captured in the SHP2^E76K^ crystal structure (Pádua *et al*, 2018). More recent studies indicate that the active state of SHP2 is better described as a dynamic, structurally heterogeneous ensemble rather than a single static conformation (Calligari *et al*, 2021; Anselmi & Hub, 2023). Consistent with this view, other activating mutations in SHP2 have also been shown to populate open states, and studies of SHP2^E76A^ revealed a broad distribution of interdomain distances, again pointing to the presence of multiple accessible open conformations (LaRochelle *et al*, 2016). Thus, while the E76K mutation relieves auto-inhibition and is therefore largely incompatible with the closed conformation, it is able to access a wide range of open states (**Figure 4A**).

**Figure 4.**
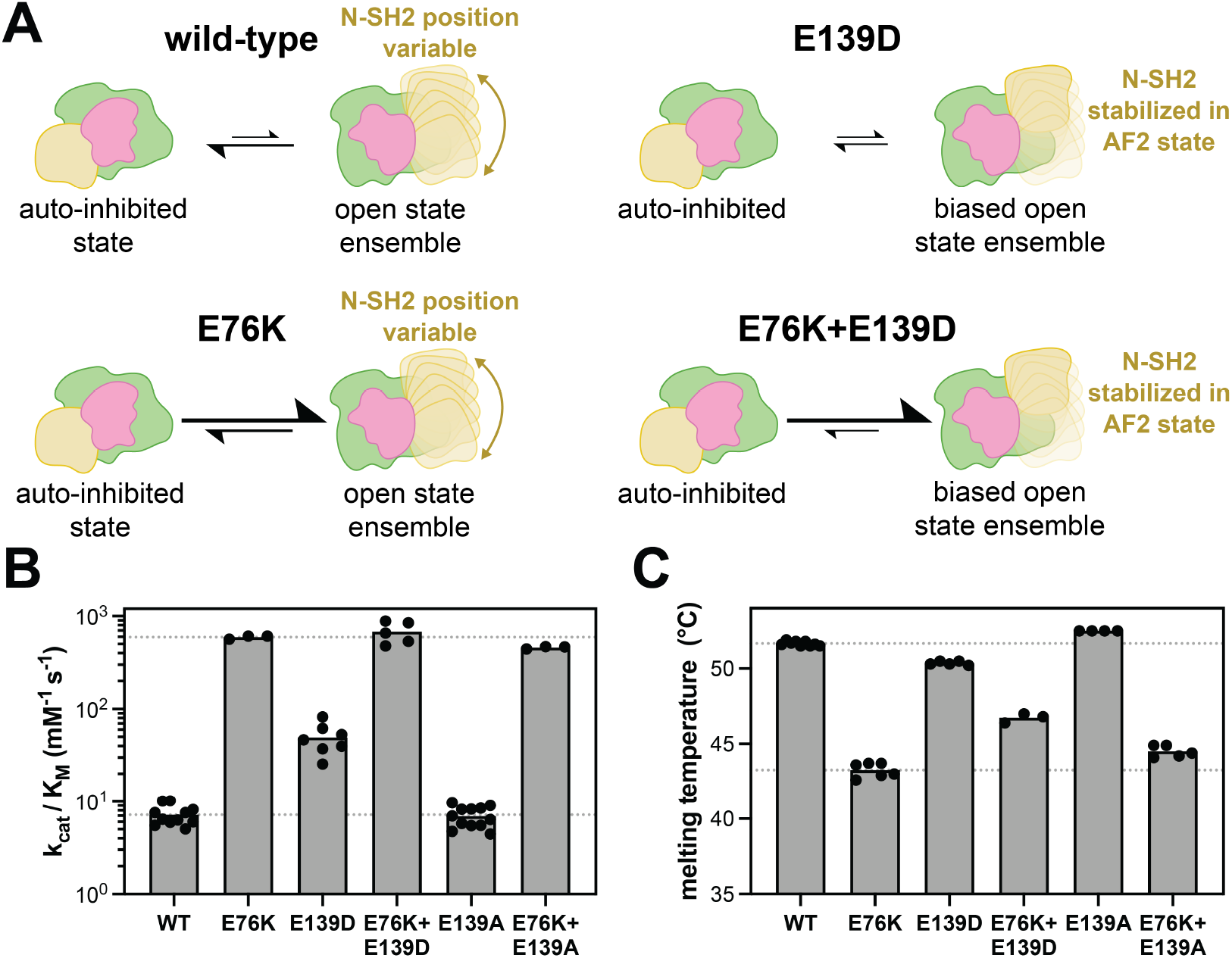
Divergent effects of E76K and E139D mutations on SHP2 activity and stability. (**A**) Cartoon schematic of the mutational effects of E76K and E139D on the SHP2 open-close equilibrium and on SHP2 structure, particularly N-SH2 positioning. Equilibrium arrows reflect mechanistic assumptions, specifically that E76K destabilizes the auto-inhibited state, whereas E139D stabilizes an open state. (**B**) Basal catalytic activity measurements comparing E76 and E139 mutations, individually and in combination. SHP2^E76K^ and SHP2^E76K+E139D^ show comparable activity, whereas SHP2^E76K+E139A^ activity is slightly diminished. Dashed lines equal SHP2^WT^ and SHP2^E76K^ activity. (**C**) Differential scanning fluorimetry measurements of E76 and E139 mutants, individually and in combination, supporting the hypothesis that the E76K mutation destabilizes the auto-inhibited state, whereas the E139D mutation stabilizes an open state. Dashed lines equal SHP2^WT^ and SHP2^E76K^ melting temperatures.

By contrast, based on our AlphaFold2 model and biochemical results, the E139D mutation appears to bias the open-state conformational ensemble of SHP2 by preferentially stabilizing a specific position and orientation of the N-SH2 domain (**Figure 4A**). However, based on the fact that the E139D mutation only increases basal activity 7-fold, and this mutation does not disrupt the auto-inhibitory interface, we hypothesize that SHP2^E139D^ still rests partly in the auto-inhibited state, unlike SHP2^E76K^. To further characterize the mechanistic differences between these two mutations, we made an E76K+E139D double mutant. Whereas the E76K mutation alone should fully destabilize the auto-inhibited state to yield an ensemble of active states, addition of the E139D mutation might bias SHP2 toward a specific open state (**Figure 4A**). First, we measured catalytic efficiencies for both single mutants and the double mutant (**Figure 4B**). While both single mutants increase catalytic activity, as expected, SHP2^E76K^ is demonstrably much more active against the DiFMUP substrate than SHP2^E139D^. The SHP2^E76K+E139D^ double mutant was comparable to SHP2^E76K^ in terms of catalytic activity (**Figure 4B**). This suggests that the conformational state stabilized by E139D can achieve maximal SHP2 catalytic activity, provided the auto-inhibited state cannot be accessed. As a control, we also made the E76K+E139A double mutant. We found that the activity of SHP2^E76K+E139A^ was slightly lower than that of SHP2^E76K^ alone (**Figure 4B**, p-value < 0.01), but still substantially higher than all variants lacking the E76K mutation, suggesting that Glu139 may play a modest role in stabilizing an active state of SHP2.

Next, we performed differential scanning fluorimetry to measure the thermal stability of SHP2 and make inferences about its conformations as a function of mutations (**Supplementary Table 2**). Previous work has shown that melting temperature (T_m_) can indirectly report on SHP2 conformation (Serbina & Bishop, 2023; Kim *et al*, 2024; Jiang *et al*, 2025). Specifically, mutants in which the auto-inhibited state is destabilized (e.g. SHP2^E76K^) generally have lower T_m_ values than SHP2^WT^, most likely due to fewer interdomain contacts. By extension, given that SHP2^E139D^ stabilizes a specific open conformation with new interdomain contacts, we hypothesized that it may have a higher T_m_ value than SHP2^E76K^. Thus, we measured the melting temperatures for both single mutants and the SHP2^E76K+E139D^ double mutant (**Figure 4C**). Consistent with previous work, we observed a large decrease in melting temperature for SHP2^E76K^ when compared to SHP2^WT^ (Serbina & Bishop, 2023; van Vlimmeren *et al*, 2024; Jiang *et al*, 2025). By comparison, SHP2^E139D^ only showed a slight decrease in T_m_ relative to SHP2^WT^, however interpretation of the magnitude of this decrease is complicated by the fact that the SHP2^E139D^ likely still partly occupies the auto-inhibited state. Notably, the SHP2^E76K+E139D^ double mutant had a higher T_m_ than SHP2^E76K^, despite the fact that both variants should not appreciably adopt the auto-inhibited state (**Figure 4C**). This is consistent with the idea that the E139D mutation enhances specific interdomain contacts in an active/open conformation, which could elevate thermal stability, whereas E76K merely destabilizes the auto-inhibited state (**Figure 4A**).

Interestingly, constructs bearing the SHP2^E139A^ mutation had slightly elevated melting temperatures relative to their Glu139 counterparts (**Figure 4C**). One speculative explanation for this is that both Glu139 and Asp139 can participate in AF2 open-state stabilizing interactions with R5 (and potentially R4), whereas A139 cannot. Consequently, the E139A mutation may lead to a slightly higher population of auto-inhibited SHP2 by destabilizing the AF2 open state. Indeed, E139A augments T_m_ in every context except for the R4A/R5A background, where there is no ion pairing partner available in the AF2 state. We note that the effect of the E139A mutation is not reflected in activity measurements of SHP2^WT^ and SHP2^E139A^, which have nearly identical but low catalytic efficiencies (**Figure 4B**). However, the E139A mutation does diminish activity in the presence of E76K (**Figure 4B**), suggesting that the putative destabilization of the AF2 open state by E139A is more detectable when the protein has higher occupancy of open states to begin with. Finally, we note that not all of our thermal stability data can be easily explained using our current structural models and biochemical data. For example, the E139D mutation lowers the T_m_ of SHP2 even in an R4A/R5A background (**Supplementary Table 2**), despite this mutation having no effect on activity in this context. These observations underscore the challenges in interpreting thermal denaturation curves on a multi-conformational ensemble for a multi-domain protein, where not all conformations may be structurally resolved.

### The E139D mutation alters the phosphoprotein binding preferences of SHP2

SHP2 is known to tightly bind bis-phosphorylate peptides and proteins through simultaneous engagement of the N-SH2 and C-SH2 domains with nearby phosphosites (Myers *et al*, 1998; Cunnick *et al*, 2001; Keilhack *et al*, 2005). Indeed, bis-phosphorylated molecules are some of the most potent activators of SHP2 and play a key role in coordinating SHP2 signaling (Hayashi *et al*, 2017; Marasco *et al*, 2020). Given that the E139D mutation appears to bias the SHP2 conformational ensemble toward a specific open state, we reasoned that this might narrow the range of possible SH2 orientations relative to one another, thereby altering binding preferences for bis-phosphorylated proteins. To test this, we first analyzed our MD trajectories. We measured the dihedral angles between two planes defined by the center of mass of either SH2 domain (Cα of L43 and L149) and the Cα of Arg111 and Arg220 on the linkers (**Supplementary Figure 6A**). This dihedral angle is consistently sharper in the SHP2^WT/E139D^/AF2 simulations when compared to the SHP2^WT^/6CRF simulations, correlating with the further rotation of the N-SH2 domain relative to the C-SH2 domain (**Supplementary Figure 6B**). This repositioning of SH2 domains can also alter the orientation of the phosphotyrosine binding residues in the N- and C-SH2 domains. We measured the distance between the Cα atoms of the two phosphotyrosine binding arginine residues in the SH2 domains, Arg32 and Arg138, in both the 6CRF and the AF2 simulations (**Figure 5A**). The arginine residues in the N- and C-SH2 domain start off at similar distances in the AF2 model and SHP2^E76K^ crystal structure, but diverge during simulations, with the AF2-derived simulations maintaining shorter distances than the 6CRF-derived simulations (**Figure 5B**). Moreover, the AF2 simulations show a tighter range of distances between Arg32 and Arg138. We note that these distances represent a direct path from Arg32 to Arg138, as a proxy for inter-pocket distances, and the actual path that a bis-phosphorylated polypeptide would have to take to simultaneously bind both domains without steric clashes would be longer.

**Figure 5.**
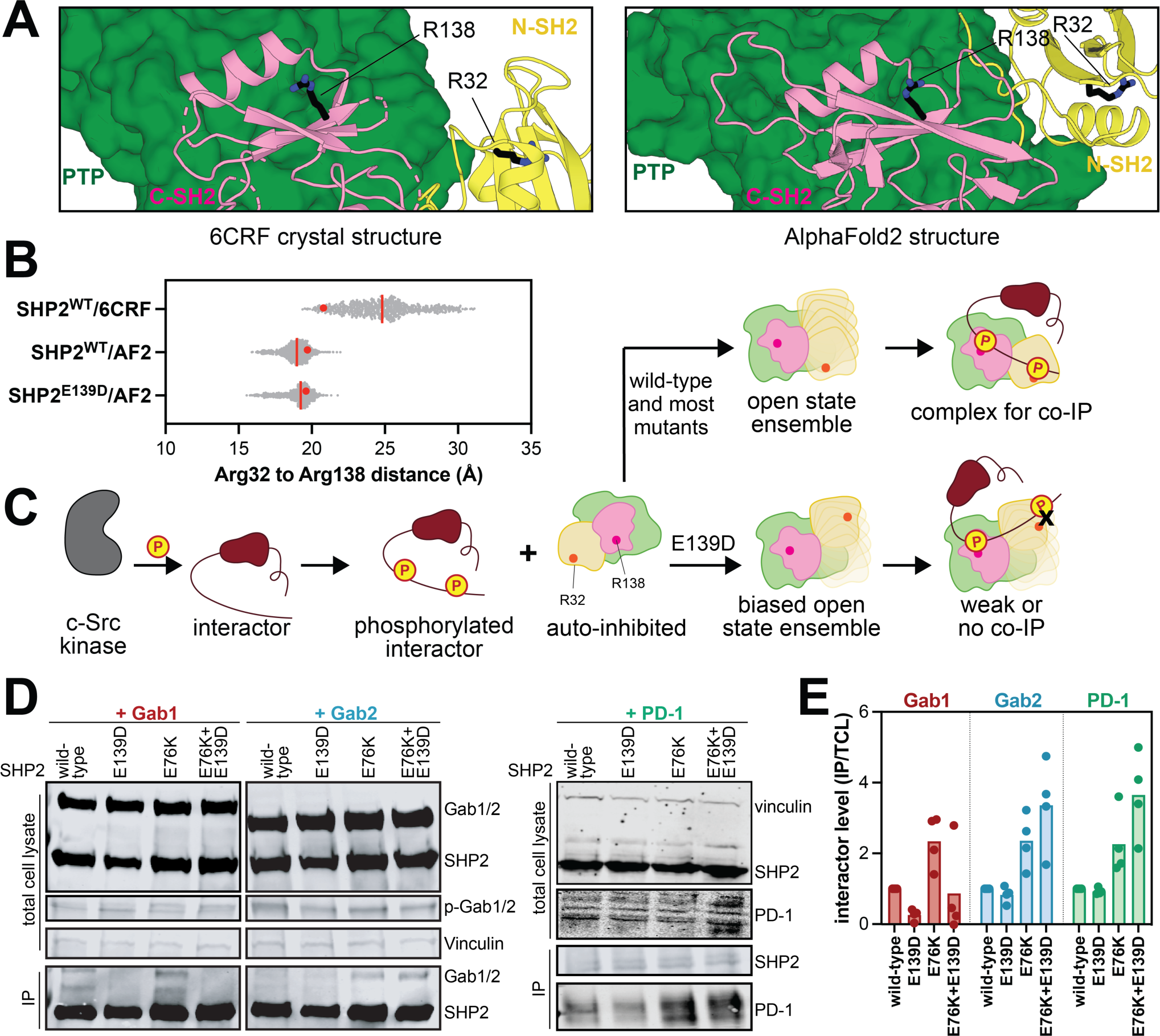
The E139D mutation alters the scope of SHP2-phosphoprotein interactions. (**A**) Positions of Arg32 and Arg138 in 6CRF and AF2 structures. (**B**) Distances (Å) between Arg32 and Arg138 Cα atoms in SHP2^WT^/6CRF, SHP2^WT^/AF2, and SHP2^E139D^/AF2 simulations. The red line indicates the median value of the distribution, and the red dot denotes the measurement in the starting model used for that simulation. (**C**) Schematic depicting how the E139D-stabilized open state might impact phosphoprotein binding. (**D**) Representative western blots showing altered co-immunopurification trends for Gab1, Gab2, and PD-1 by the E139D mutation. (**E**) Quantification of SHP2 co-immunopurification results for Gab1, Gab2, and PD-1 based on 4 biological replicates for each interacting protein. Interactor levels are normalized to Gab1/Gab2/PD-1 protein levels in the total cell lysate (TCL), then further normalized to Gab1/Gab2/PD-1 levels immunopurified in the wild-type sample.

To examine whether the geometric constraints suggested by the AF2 model and our MD simulations could impact SHP2 interactions, we assessed the ability of different SHP2 variants to bind to three different bis-phosphorylated proteins that are known SHP2 interactors, Gab1, Gab2, and PD-1. We reasoned that, if SHP2^E139D^ merely adopts any more open conformation, this would enhance the pull-down of the interacting phosphoprotein, as we have previously shown for SHP2^E76K^ (van Vlimmeren *et al*, 2024). However, if SHP2^E139D^ is biased toward a conformation that is incompatible with the distance between the phosphosites on the interacting protein, we would expect to see reduced pull-down of that protein when compared to SHP2^WT^ or SHP2^E76K^ (**Figure 5C**). Thus, we co-expressed SHP2 with Gab1, Gab2, or PD-1 in HEK 293 cells. A constitutively active variant of c-Src was also co-expressed to ensure Gab1/Gab2/PD-1 phosphorylation (van Vlimmeren *et al*, 2024). SHP2 was immuno-purified, and levels of co-immunopurified activator protein were assessed by western blot. Interestingly, compared to SHP2^WT,^ we saw a reduction of Gab1 binding for SHP2^E139D^, whereas Gab2 and PD-1 binding was similar between the two SHP2 variants (**Figure 5D,E and Supplementary Figure 7**). By contrast, SHP2^E76K^ showed increased immunopurification for all three proteins, as expected. Finally, we also made a double mutant of SHP2^E76K+E139D^ to eliminate any effect of the auto-inhibited state and instead only consider the effect of E139D in stabilizing a specific open conformation. Addition of the E139D mutation on top of the E76K mutation reduced binding to Gab1 when compared to SHP2^E76K^, consistent with the conformationally constraining role of the E139D mutation (**Figure 5D,E**). By contrast, the SHP2^E76K+E139D^ double mutant had similar or enhanced immunopurification of Gab2 and PD-1 relative to SHP2^E76K^, suggesting that the E139D mutation stabilizes a state that is compatible with Gab2 and PD-1 binding (**Figure 5D,E**). Consistent with this observation, PD-1 has previously been suggested to bind to SHP2 in a conformation that resembles the SHP1 open-conformation crystal structure (PDB code 3PS5), which is similar to our proposed E139D-stabilized AF2 conformation (Marasco *et al*, 2020). Overall, these results support the hypothesis that the E139D mutation stabilizes a specific open conformation that can alter SHP2 ligand binding compatibility.

One alternative explanation for our co-immunopurification results is that the E139D mutation alters the intrinsic binding properties of the C-SH2 domain. Indeed, this residue is directly adjacent to R138, which is essential for phosphoprotein binding (Marasco *et al*, 2020; van Vlimmeren *et al*, 2024). A prior report suggested that this mutation shifts the binding specificity of the isolated C-SH2 to somewhat resemble that of the N-SH2 domain (Martinelli *et al*, 2008). However, we previously showed that the E139D mutation does not appreciably impact binding affinity or specificity against array of phosphopeptides, including those derived from the Gab1, Gab2, and PD-1 phosphosites (**Supplementary Figure 8A,B**) (van Vlimmeren *et al*, 2024). Thus, it is more likely that the E139D mutation alters binding preferences through a change in interdomain orientation and dynamics. In support of this model, we analyzed our previously reported proximity-labeling proteomics data for SHP2^WT^, SHP2^E76K^, and SHP2^E139D^ (van Vlimmeren *et al*, 2025). These three variants share a core interactome but also show differences in proximity labeling for a number of proteins (**Supplementary Figure 8C**). This includes a SHP2^E139D^-specific lack of significant labeling of Gab1 over the negative control (**Supplementary Figure 8D**). In our proximity-labeling proteomics datasets, we also observed an E139D-dependent enhancement in significant labeling of MPZL1, another disease-relevant SHP2 interactor (Paardekooper Overman *et al*, 2014). A recent crystal structure of the SHP2^WT^ tandem SH2 domains bound to MPZL1 showed that the SH2 domains adopt an arrangement that resembles that seen our AF2 model and the SHP1 active state structure (Perla *et al*, 2025). Notably, another study that used the SHP2 tandem SH2 domains for immunopurification and phosphoproteomics also reported different interactomes for the wild-type and E139D tandem SH2 domains (Müller *et al*, 2013). Critically, that investigation used a construct that spanned SHP2 residues 2-220 and thus had the potential to maintain stabilizing interactions between Arg4/Arg5 and Asp139. Collectively, our results show that the E139D mutation can not only impact SHP2 basal activity, but also perturb SHP2 protein-protein interaction specificity, which could contribute to the pathogenicity of this variant.

## Discussion

The first structure of SHP2 in a catalytically-competent state, determined using the E76K mutant, was a major milestone for the SHP2 field (LaRochelle *et al*, 2018). Since then, it has become evident that SHP2 can access multiple open conformations and that the SHP2^E76K^ crystal structure represents just one of the possible states (LaRochelle *et al*, 2016; Pádua *et al*, 2018; Tao *et al*, 2021; Calligari *et al*, 2021; Anselmi & Hub, 2023). Using AlphaFold2, we previously obtained a model of SHP2^WT^ in an open conformation that is distinct from the SHP2^E76K^ structure (Jiang *et al*, 2025) — one that more closely resembles the open conformation previously described for SHP1 (Wang *et al*, 2011). Notably, in the open conformation crystal structure of SHP1^C453S^, residue Glu137 (the equivalent of SHP2 Glu139) forms interactions with an arginine residue near the N-terminus, and we see a similar ion pair between Glu139 and Arg5 in our AlphaFold2 model (**Figure 1G,H**). These observations led us to probe whether such an interaction could help explain the pathogenic effect of the E139D mutation.

In this study, we analyzed MD simulations starting from our open conformation AF2 model, for both SHP2^WT^ and SHP2^E139D^, which revealed that Glu/Asp139 can form electrostatic interactions with both Arg4 and Arg5. Through biochemical measurements, we have now shown that the activating effect of the E139D mutation is significantly dampened in the background of the R5A mutation, suggesting that Arg5 promotes SHP2 activity through energetic coupling with Asp139. This observation cannot be explained by current crystallographic states of SHP2 (Hof *et al*, 1998; LaRochelle *et al*, 2018; Yu *et al*, 2013), and it and supports the AlphaFold2 model as one plausible alternative open conformation. Notably, our computational data for SHP2^WT^, combined with activity and stability measurements with SHP2^E139A^ and double mutants, suggest that the wild-type residue, Glu139, may also play a modest role in stabilizing the AF2 state. Thus, this particular conformational state, not observed in any SHP2 crystal structure, is likely to be relevant to the wild-type enzyme, in addition to the E139D mutant.

Unlike the clear roles for Glu/Asp139 and Arg5, which primarily make interactions in the AF2-modeled conformation, the role of Arg4 in controlling SHP2 structure appears multi-faceted. The MD simulations based on the AF2 open state model suggest that Arg4 plays a larger role than Arg5 in stabilizing the AF2 state via interactions with Glu/Asp139 in this conformational state. However, in our experiments, loss of Arg4 (R4A mutation) increased the basal activity of SHP2. This is likely because Arg4 forms a salt bridge with Glu258, and potentially also Glu252, in the auto-inhibited state (Hof *et al*, 1998; Yu *et al*, 2013). Arg4 also forms a salt bridge with Asp485 in the 6CRF open state (LaRochelle *et al*, 2018). Thus, it is plausible that the R4A mutation destabilizes both the auto-inhibited and 6CRF open states, thereby enhancing occupancy of the AF2 open state, which can still be stabilized by an Arg5 ion pair with Glu/Asp139.

It is important to note that the standard MD simulations presented in this study, and in our past work (Jiang *et al*, 2025), cannot be used to quantitatively dissect the contributions of each ion pair to the energetics of each SHP2 conformational state. Due to their short timescale (microseconds) relative to the actual timescale of SHP2 interdomain dynamics (seconds) (Pádua *et al*, 2018; Tao *et al*, 2021), these simulations only provide a local sampling of conformational space around each starting conformation. Several recent studies have implemented enhanced sampling methods to probe a broader swath of the SHP2 conformational and energetic landscape (Calligari *et al*, 2021; Tao *et al*, 2021; Hou *et al*, 2023). These approaches, applied to Arg4, Arg5, and E139 mutants individually and in combination, are likely to provide further insights into the importance of the AF2 open state and the effect of the E139D mutation.

One consequence of the E139D mutation stabilizing the AF2 active state is that it constrains SHP2 to adopt a narrower range of distances between the two phosphotyrosine-binding pockets in its tandem SH2 domains. It is plausible that this would bias SHP2 to interact more tightly with bis-phosphorylated proteins that are compatible with this conformational constraint, while weakening binding to those bis-phosphorylated proteins that cannot conform to this interdomain arrangement. Binding experiments with three known bis-phosphorylated SHP2 interactors, Gab1, Gab2, and PD-1, support this idea by showing that the E139D mutation selectively reduces binding to Gab1. This is in contrast to mutations like E76K, which activate SHP2 by destabilizing the active state but do not bias the open state ensemble toward any one particular conformation. Ultimately, a deeper understanding of this change in binding specificity awaits the determination of high-resolution structures of full-length or near-full-length of SHP2 bound to different bis-phosphorylated polypeptides. Regardless, an important consequence of the biased binding seen for SHP2^E139D^ is that this mutant may be activated by a different spectrum of upstream phosphoproteins than SHP2^WT^ and other SHP2 mutants. This biased activation, however, should only be relevant to bis-phosphorylated but not mono-phosphorylated proteins. Indeed, past studies have shown the E139D mutation marginally impacts the EC_50_ for activation by a mono-phosphorylated peptide, whereas some bis-phosphorylated peptides dramatically enhance activation of SHP2^E139D^ when compared with SHP2^WT^ (Keilhack *et al*, 2005; Tartaglia *et al*, 2006; Martinelli *et al*, 2008). The downstream signaling outcomes associated with this mutation-dependent rewiring of SHP2 interactions will require further investigation.

In summary, our results suggest that Glu139 plays a critical role in stabilizing a novel active state of SHP2 through interactions with Arg5 and potentially Arg4. Substitution of Glu to Asp at this position strengthens these interactions, providing a mechanistic explanation for the activating effect of the disease associated E139D mutation. More broadly, this work underscores the importance of considering protein conformational dynamics and uncharacterized protein states when interpreting the functional consequences of disease-associated mutations.

## Acknowledgements

We would like to thank the members of the Shah for their scientific insights and helpful discussions. This research was funded by NIH/NIGMS grant R35GM138014 to NHS. This work used the Expanse GPU cluster at the San Diego Supercomputer Center through allocation BIO220139 from the Advanced Cyberinfrastructure Coordination Ecosystem: Services & Support (ACCESS) program, which is supported by NSF grants 2138259, 2138286, 2138307, 2137603, and 2138296.

## Author Contributions

A.E.v.V and Z.J. contributed equally to this work. A.E.v.V, Z.J., and N.H.S. conceived the project, designed the experiments, and wrote the manuscript. A.E.v.V., Z.J., and A.T.A.L. conducted the experiments. D.K. conducted the molecular dynamics simulations. A.E.v.V., Z.J., D.K., A.T.A.L., and N.H.S. analyzed and interpreted the data and edited the manuscript.

## Data and materials availability

Molecular dynamics trajectory files for SHP2^WT^/6CRF and SHP2^WT^/AF2 are available via the following Dryad repository: https://doi.org/10.5061/dryad.83bk3jb18. Molecular dynamics trajectory files for SHP2^E139D^/AF2 are available via the following Dryad repository: https://doi.org/10.5061/dryad.0rxwdbscv. All other datasets associated with this work are available as Supplementary Information.

## Supplementary Information

### Supplementary Information file contains

Supp. Figure 1. Unique activating effect of the E139D mutation. Supp. Figure 2. Measurements of SH2 domain position and rotation.

Supp. Figure 3. Solvent accessible surface area of Glu/Asp139 in MD simulations. Supp. Figure 4. Mutant cycle analyses with R4A, R5A, and E139D.

Supp. Figure 5. Mutant cycle analyses with R4A, R5A, and E/D139A.

Supp. Figure 6. Interdomain dihedral angle analysis showing SH2 domain repositioning. Supp. Figure 7. Uncropped blots for SHP2-phosphoprotein co-immunopurification.

Supp. Figure 8. Comparison of peptide binding and proximity-labeling proteomics for SHP2 variants. Materials and methods

Supplementary References

### Supplementary tables can be found as separate spreadsheet files

Supp. Table 1. Catalytic parameters for all SHP2 variants used in this study. Supp. Table 2. Melting temperatures of all SHP2 variants used in this study.

