## Supplementary Figures and Methods for "The pathogenic E139D mutation stabilizes a non-canonical active state of the multi-domain phosphatase SHP2"

[3] Independent researcher

[4] Herbert Irving Comprehensive Cancer Center, Columbia University, New York, NY 10032

[†] these authors contributed equally

#### Table of contents:

| | | |
|---|---|---|
| Supp. Figure 1 | Unique activating effect of the E139D mutation. | pg 2 |
| Supp. Figure 2 | Measurements of SH2 domain position and rotation. | pg 3 |
| Supp. Figure 3 | Solvent accessible surface area of Glu/Asp139 in MD simulations. | pg 4 |
| Supp. Figure 4 | Mutant cycle analyses with R4A, R5A, and E139D. | pg 5 |
| Supp. Figure 5 | Mutant cycle analyses with R4A, R5A, and E/D139A. | pg 6 |
| Supp. Figure 6 | Interdomain dihedral angle analysis showing SH2 domain repositioning. | pg 7 |
| Supp. Figure 7 | Uncropped blots for SHP2-phosphoprotein co-immunopurification. | pg 8 |
| Supp. Figure 8 | Comparison of proximity-labeling hits for WT, E76K, and E139D. | pg 9 |
| Materials and methods | | pg 10 |
| Supplementary references | | pg 13 |

**Supplementary Tables** (included as separate spreadsheet files):
Supp. Table 1. Catalytic efficiencies of all SHP2 variants used in this study.
Supp. Table 2. Melting temperatures of all SHP2 variants used in this study.

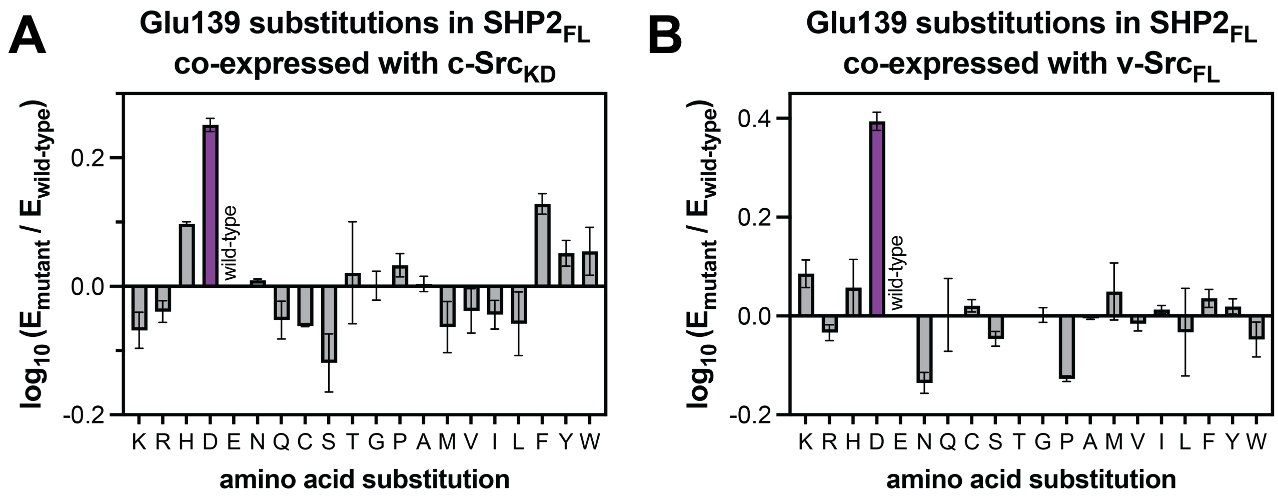


**Supplementary Figure 1. Unique activating effect of the E139D mutation.** Mutational effects at Glu139 were extracted from a deep mutational scanning study of SHP2 in a yeast selection assay (Jiang *et al*, 2025). (**A**) Mutational effects at Glu139 in the context of full-length SHP2 co-expressed with the c-Src kinase domain (weak selective pressure). (**B**) Same as (**A**), but with full-length SHP2 and full-length v-Src (strong selective pressure).

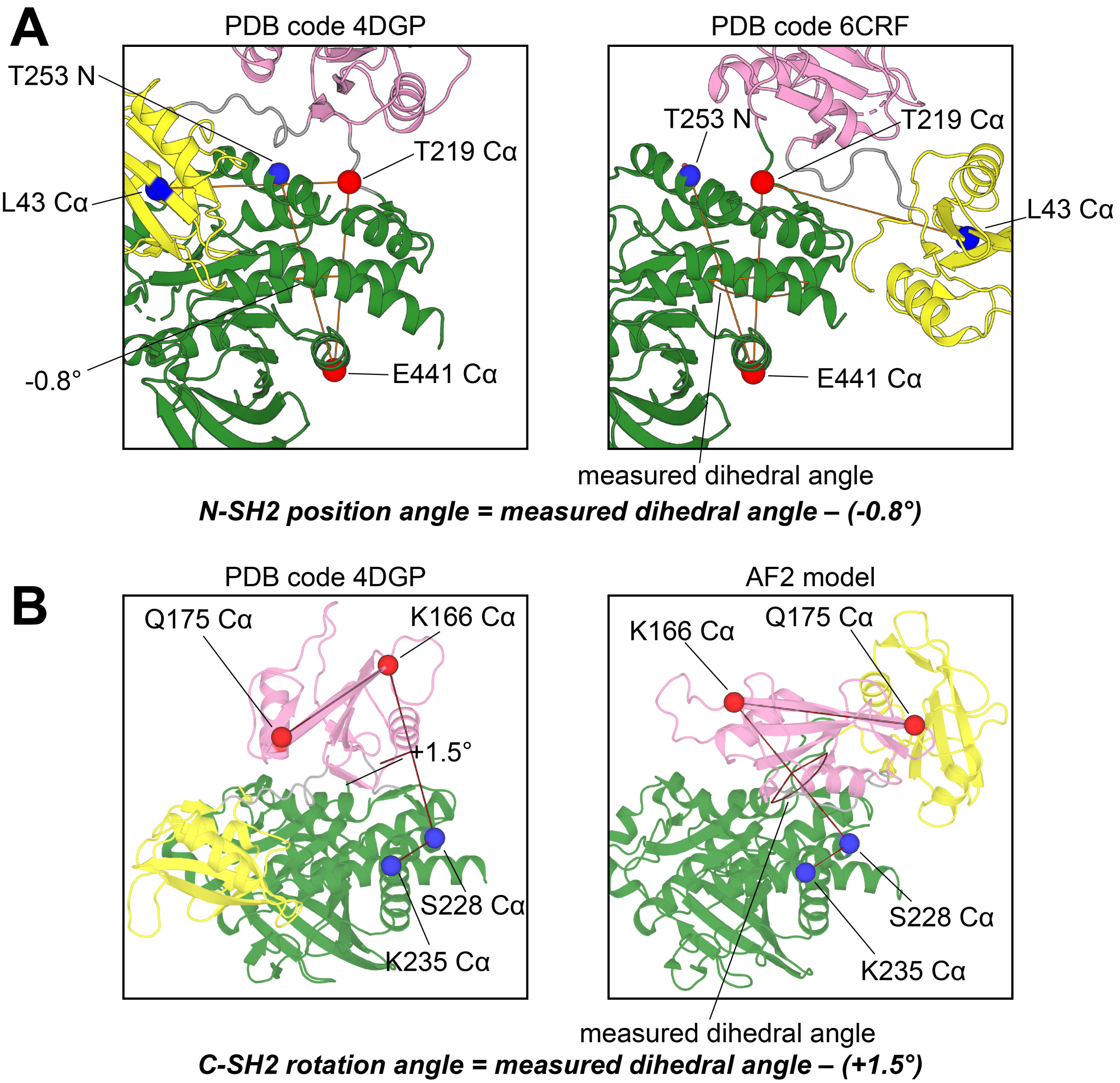


**Supplementary Figure 2. Measurements of SH2 domain position and rotation.** (**A**) To measure N-SH2 position, we first identified an atom that approximates its center of mass, the Cα atom of L43. Then, we defined an axis of rotation by T219 Cα and E441 Cα in the PTP domain. In the autoinhibited state (PDB code 4DGP), the vector defined by L43 Cα and T219 Cα nearly passes through the main chain nitrogen atom of T253, which is relatively static in our MD trajectories and between different conformational states. Thus, T253 N is a good proxy for the position of the auto-inhibited state N-SH2 domain. Indeed, the dihedral angle defined by L43 Cα, T219 Cα, E441 Cα, and T253 N is -0.8°. We measured this dihedral angle for each frame in our open conformation MD trajectories, then subtracted -0.8° to determine N-SH2 domain positioning relative to the auto-inhibited state. (**B**) To measure rotation of the C-SH2 domain on the face of the PTP domain, we used the vector defined by K116 Cα and Q175 Cα to represent C-SH2 orientation. The vector defined by S228 Cα and K235 Cα in the PTP domain is a good proxy for C-SH2 domain orientation in the auto-inhibited state, as it is nearly parallel to the C-SH2 vector. Indeed, the dihedral angle defined by K235 Cα, S228 Cα, K116 Cα, and Q175 Cα is +1.5° in the auto-inhibited state. We measured this dihedral angle for each frame in our open conformation MD trajectories, then subtracted +1.5° to determine C-SH2 rotation relative to the auto-inhibited state.

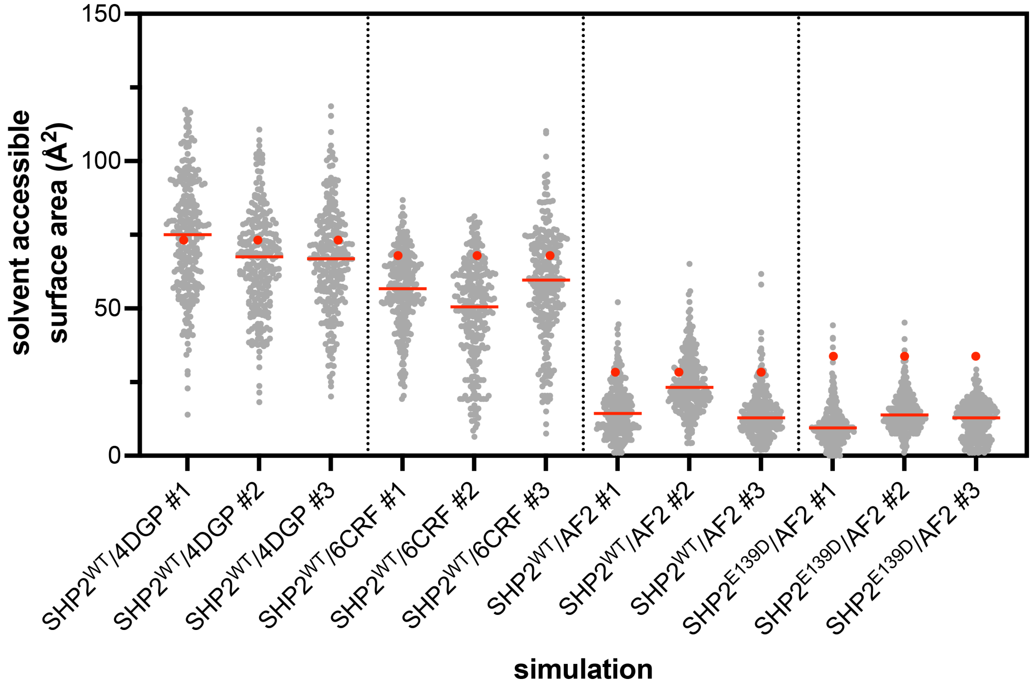


**Supplementary Figure 3. Solvent accessible surface area of Glu/Asp139 in MD simulations.** Data from triplicate 2.5 μs simulations in various states are shown, highlighting a more buried Glu/Asp139 in simulations based on the AF2 model. The red line indicates the median value of the distribution, and the red dot denotes the measurement in the starting model used for that simulation.

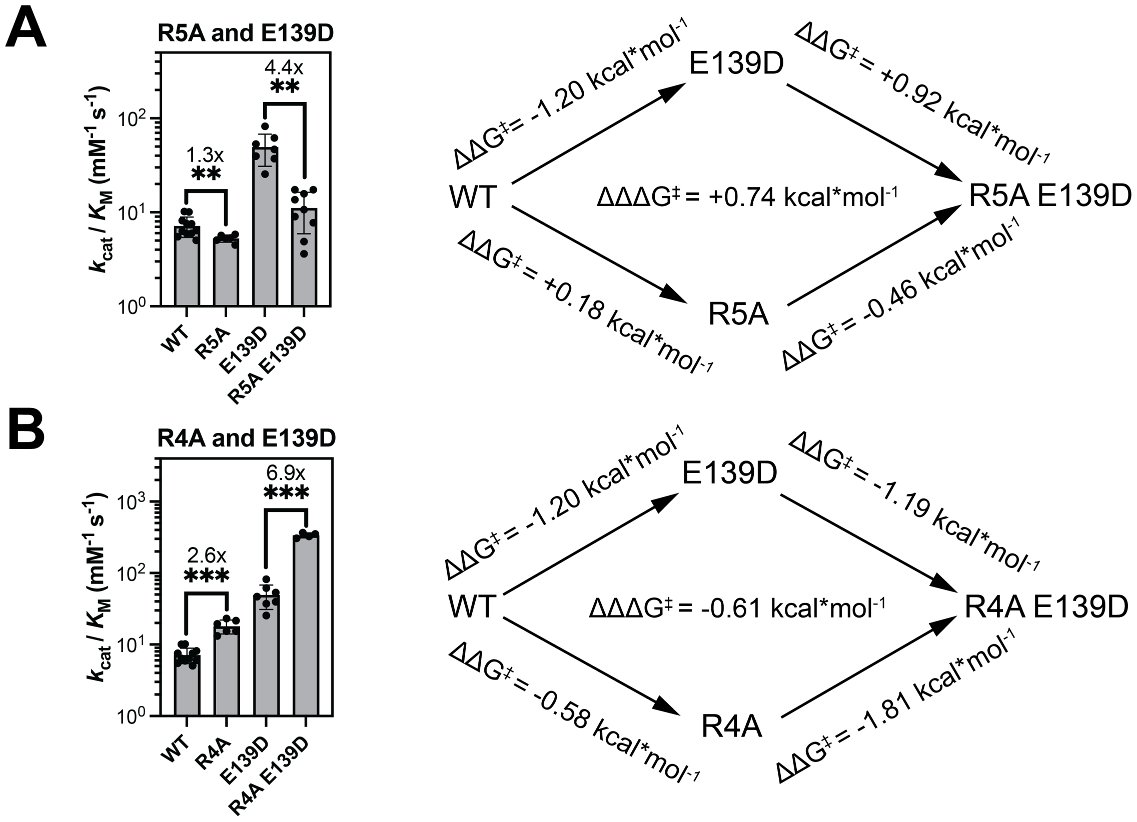


**Supplementary Figure 4. Mutant cycle analyses with R4A, R5A, and E139D.** Double mutant cycles of (**A**) R5A and E139D and (**B**) R4A and E139D. Bar graphs represent an alternative comparison from the ones described in **Figure 3B,C**. On the right are depictions of the mutant cycles, showing the $\Delta\Delta G^{\ddagger}$ and coupling energy $\Delta\Delta\Delta G^{\ddagger}$ calculations. In all panels, statistical significance was assessed by Welch's two-tailed t test. ns = not significant, * denotes $p < 0.05$, ** denotes $p < 0.01$, *** denotes $p < 0.001$.

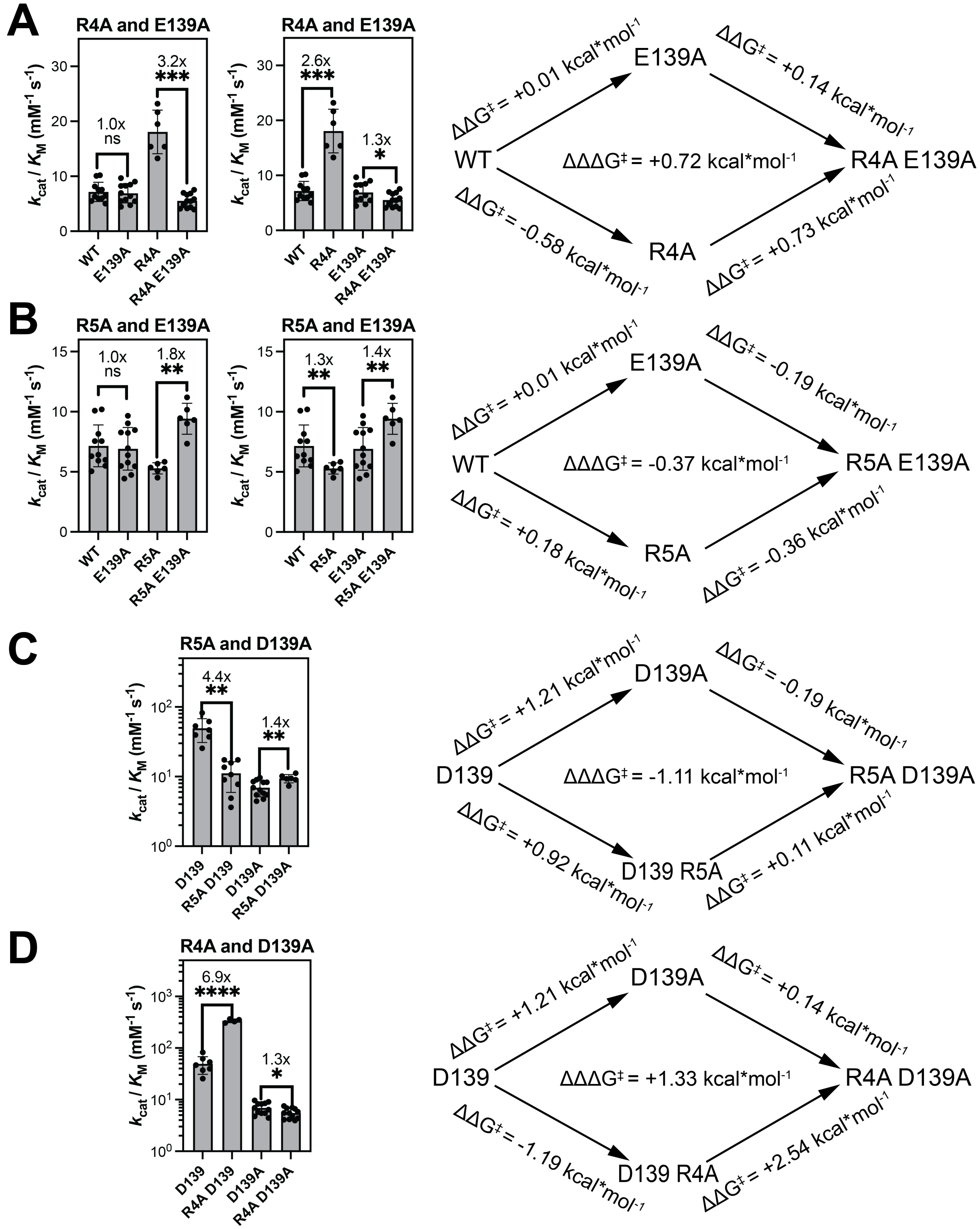


**Supplementary Figure 5. Mutant cycle analyses with R4A, R5A, and E/D139A.** Double mutant cycles of (**A**) R4A and E139A, (**B**) R5A and E139A, (**C**) R4A and D139A, and (**D**) R5A and D139A. For panels (**C**) and (**D**), bar graphs represent an alternative comparison from the ones described in **Figure 3F,G**. On the right are depictions of the mutant cycles, showing the $\Delta\Delta G^{\ddagger}$ and coupling energy $\Delta\Delta\Delta G^{\ddagger}$ calculations. In all panels, statistical significance was assessed by Welch's two-tailed t test. ns = not significant, * denotes $p < 0.05$, ** denotes $p < 0.01$, *** denotes $p < 0.001$.

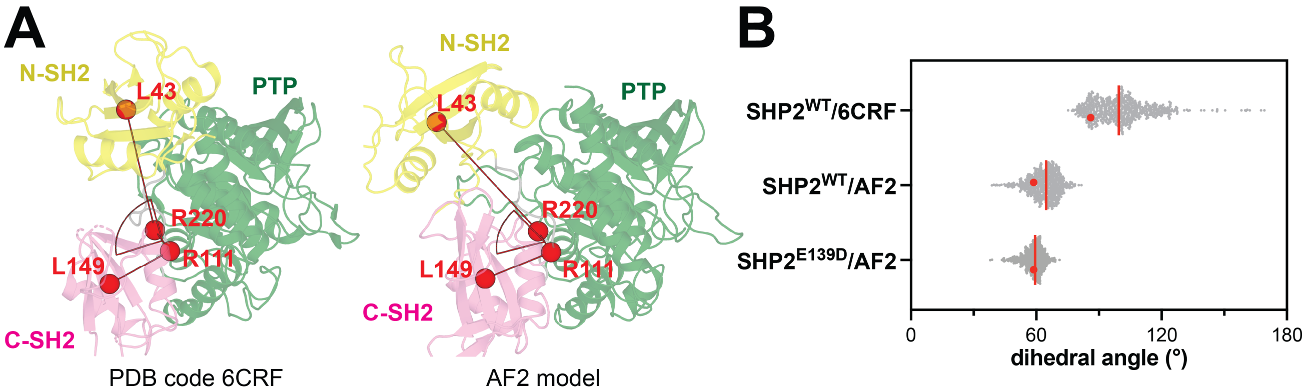


**Supplementary Figure 6. Interdomain dihedral angle analysis showing SH2 domain repositioning.** (**A**) Dihedral angles measured for quantification of the change in SH2 position. (**B**) SH2 dihedral angles in the SHP2 simulations shows sharper angle and tighter range in SHP2$^{WT}$/AF2 and SHP2$^{E139D}$/AF2 simulations when compared to the SHP2$^{WT}$/6CRF simulations. The red line indicates the median value of the distribution, and the red dot denotes the measurement in the starting model used for that simulation.

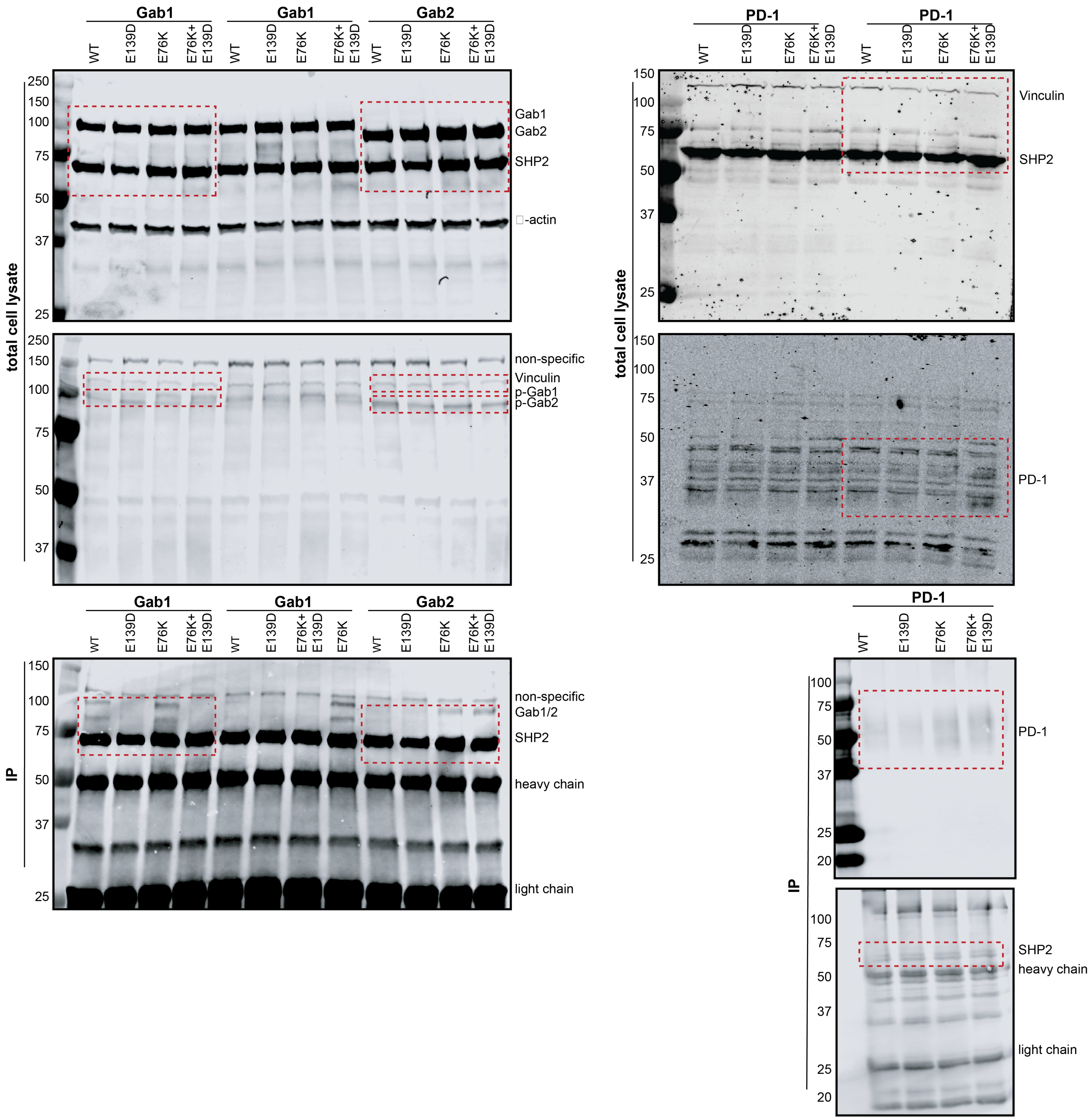


**Supplementary Figure 7. Uncropped blots for SHP2-phosphoprotein co-immunopurification.** The blots shown here correspond to the cropped blots shown in Figure 5D. The dashed red boxes correspond to the cropped regions shown in the main figure. These blots were the source of one of the four replicate datapoints in Figure 5E.

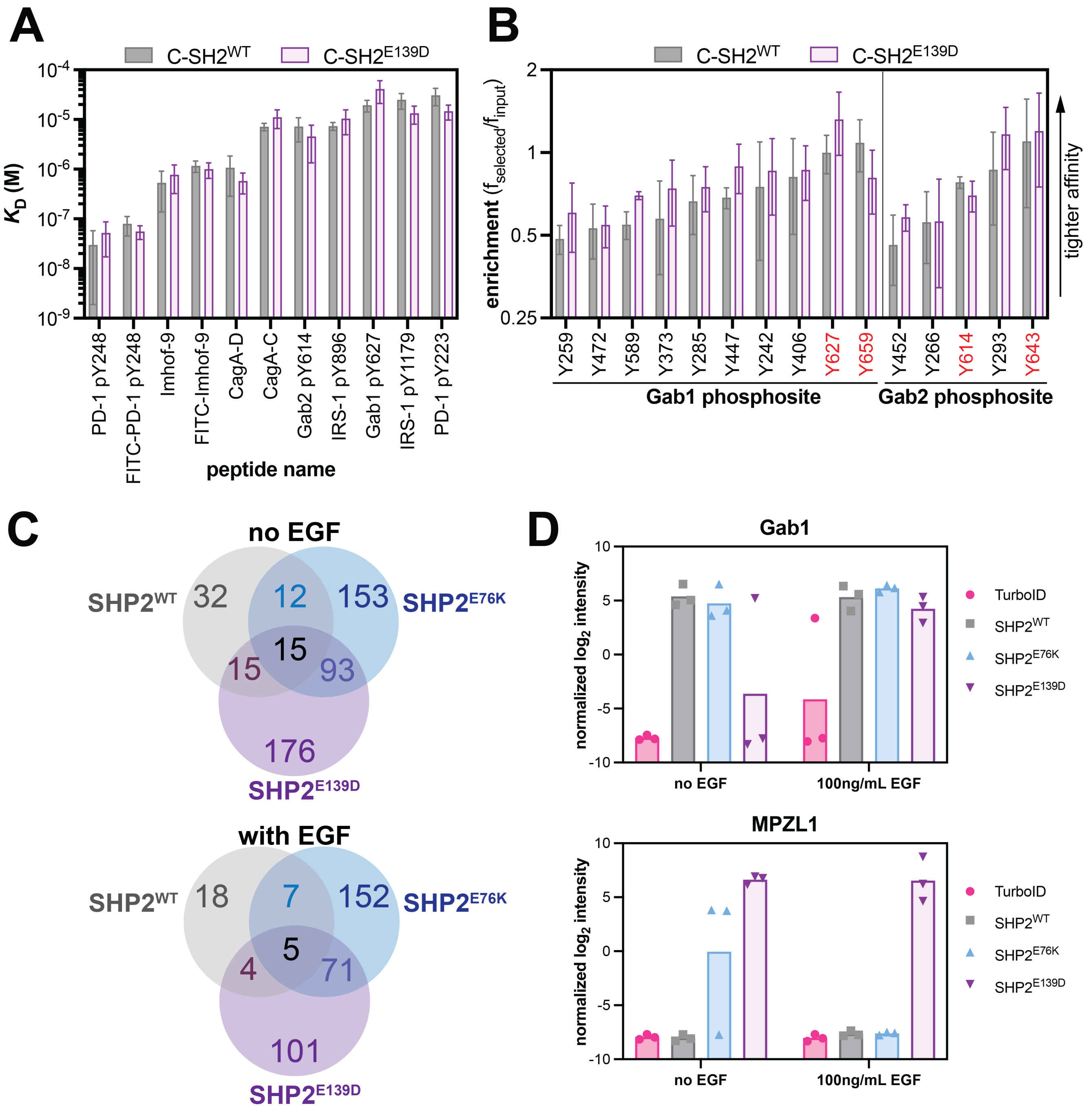


**Supplementary Figure 8. Comparison of peptide binding and proximity-labeling proteomics for SHP2 variants.** (**A**) Binding affinity measurements for various SHP2-relevant phosphopeptides with the isolated C-SH2$^{WT}$ and C-SH2$^{E130D}$ domains, measured by fluorescence polarization. N = 3-6 independent titrations. (**B**) Enrichment scores from peptide display screens phosphopeptides with the isolated C-SH2$^{WT}$ and C-SH2$^{E130D}$ domains for phosphopeptides derived from tyrosine phosphorylation sites in Gab1 and Gab2. A larger enrichment means tighter binding. The canonical SHP2 binding sites are labeled in red. Data in panels (**A**) and (**B**) are derived from a previous report on SHP2 SH2 binding specificity (van Vlimmeren *et al*, 2024). (**C**) Overlap in proximity-labeling hits that show statistically significant enrichment for SHP2-TurboID variants over a TurboID negative control, both unstimulated cells and for cells stimulated with EGF. (**D**) Normalized intensities for proximity labeling of Gab1 and MPZL1 by the TurboID-only control, as well as TurboID fusions to SHP2$^{WT}$, SHP2$^{E76K}$, and SHP2$^{E139D}$. Data in panels (**C**) and (**D**) are derived from a previous report on SHP2 proximity-labeling proteomics (van Vlimmeren *et al*, 2025).

### Materials and Methods

#### DNA constructs

The SHP2 gene was cloned from the pGEX-4TI SHP2 WT plasmid from Ben Neel (Addgene, #8322). All SHP2 genes for mammalian expression were cloned into a pEF vector with an N-terminal myc-tag. The mouse c-Src gene was expressed from the pCMV5 mouse Src plasmid, a gift from Joan Brugge and Peter Howley (Addgene, #13663). The C-terminal regulatory tail (residues 528-535) was deleted to make a constitutively active c-Src construct. The PD-1 gene was cloned from the PD-1-miSFIT-4x plasmid, which was a gift from Tudor Fulga (Addgene, #124678). The mouse Gab1 and Gab2 genes were cloned from the FRB-GFP-Gab2(Y604F/Y634F) and FRB-GFP-Gab1(Y628F/Y660F) plasmids, which were a gift from Andrei Karginov (Addgene, #188658 and #188659 respectively).

#### Molecular dynamics simulations

*Preparation of Structural Models for Simulations*

In a previous study, we conducted molecular dynamics simulations of near full-length wild-type SHP2 (residues 1-528) in three distinct conformational states: the auto-inhibited state represented by PDB code 4DGP ($SHP2^{WT}$/4DGP), an active conformation represented by PDB code 6CRF ($SHP2^{WT}$/6CRF), and a second active conformation generated by AlphaFold2 ($SHP2^{WT}$/AF2) (Jiang *et al*, 2025). The trajectories from that study were used for the analyses in this report. For this study, we also conducted simulations of the E139D mutant in the AlphaFold2 conformation ($SHP2^{E139D}$/AF2). To set up this system, we used the $SHP2^{WT}$/AF2 starting structure and mutated Glu139 to Asp using PyMOL (Schrödinger & Delano, 2020). In our original study with wild-type SHP2 simulations, crystalline waters that were present in 6CRF were added to the AlphaFold2 structure. The same approach was taken for the $SHP2^{E139D}$/AF2 simulation. Cys 459 is deprotonated in all systems, based on prior reports of the catalytic Cys of protein tyrosine phosphatases having a low pKa (Zhang & Dixon, 1993; Lohse *et al*, 1997), the observation that a C-to-S mutation in PTP1B distorts its active site (likely due to changes in electrostatics) (Scapin *et al*, 2001), and the observation that C459S disrupts SHP2 autoinhibition, whereas C459E does not (Pádua *et al*, 2018; Sha *et al*, 2023). Both N- and C-termini were capped with acetyl and amide groups, respectively, in all systems.

*Simulation Protocol*

Each system was solvated with TIP3P water (Jorgensen *et al*, 1983), and ions were added such that the final ionic strength of the system was 100 mM using the tleap package in AmberTools22 (Case *et al*, 2022). The energy of each system was minimized first for 5000 steps while holding the protein atoms and crystalline waters fixed, followed by minimization for 5000 steps while allowing all the atoms to move. Following minimization, three individual trajectories were generated for each system, with distinct initial velocities for each. The temperature of each system was raised in two stages – first to 100 K over 0.5 ns and then to 300 K over 0.5 ns. The protein atoms and crystalline waters were held fixed during the heating stage. Each system was then equilibrated for 2 ns, followed by production runs. Three production trajectories, each 2.5 µs long, were generated for each system. All equilibration runs and production runs were performed at constant temperature (300 K) and pressure (1 bar). The simulations were carried out with the Amber package (Case *et al*, 2005) using the ff14SB force field for proteins (Maier *et al*, 2015) using an integration timestep of 2 fs. The Particle Mesh Ewald approximation was used to calculate long-range electrostatic energies (Darden *et al*, 1993). All hydrogens bonded to heavy atoms were constrained with the SHAKE algorithm (Ryckaert *et al*, 1977). The Langevin thermostat was used to control the temperature with a collision frequency of 1 $ps^{-1}$. Pressure was controlled while maintaining periodic boundary conditions.

*Analyses*

MD trajectories were compiled from the raw data using the CPPTRAJ module of AmberTools22 (Case *et al*, 2022). Structures were extracted from the trajectories in both 1 ns and 10 ns increments for analysis and visualization. All measurements and calculations were done using the PDB module in

Biopython (Cock *et al*, 2009). For all calculations reported in this study, trajectories were sampled every 10 ns. The measurements from all three replicates of each system were combined to determine the reported distributions, unless otherwise stated. In cases where distance calculations involved a redundant atom (e.g. distances between two possible nitrogens and two possible oxygens in a Glu/Arg ion pair), all combinations of distance measurements were calculated, then the shortest distance at each frame was determined and used for the distribution plots. The specific atoms/residues used for measuring for N-SH2 positioning, C-SH2 rotation, and N-to-C-SH2 interdomain dihedral angle are specified in the relevant figures and legends. For visualization, trajectories were sampled every 10 ns. All structure visualization and rendering in this study was done using PyMOL (Schrödinger & Delano, 2020).

*Acknowledgements for computational resources*

This work used the Expanse GPU cluster at the San Diego Supercomputer Center through allocation BIO220139 from the Advanced Cyberinfrastructure Coordination Ecosystem: Services & Support (ACCESS) program, which is supported by an NSF grants #2138259, #2138286, #2138307, #2137603, and #2138296.

### Purification of full-length SHP2

Full-length SHP2 variants were cloned into a pET28-His-TEV plasmid. BL21(DE3) cells were transformed and grown in Terrific Broth supplemented with 100 µg/mL kanamycin at 37 °C until cells reached an OD600 of 0.5. IPTG (1 mM) was added to induce protein expression, which was performed at 18 °C overnight. Cells were centrifuged and subsequently resuspended in lysis buffer (50 mM Tris pH 7.5, 300 mM NaCl, 20 mM imidazole, 10% glycerol, and freshly added 2 mM β-mercaptoethanol). The cells were lysed using sonication (Fisherbrand Sonic Dismembrator) and spun down at 14,000 rpm for 45-60 minutes. The supernatant was applied to a 5 mL Ni-NTA column (Cytiva). The resin was washed with 10 column volumes lysis buffer and wash buffer (50 mM Tris pH 7.5, 50 mM NaCl, 20 mM imidazole, 10% glycerol, and freshly added 2 mM β-mercaptoethanol). The protein was eluted off the Ni-NTA column in elution buffer (50 mM Tris pH 7.5, 50 mM NaCl, 500 mM imidazole, 10% glycerol, and freshly added 2 mM β-mercaptoethanol) and brought onto a 5mL HiTrap Q Anion exchange column (Cytiva). The column was washed using Anion A buffer (50 mM Tris pH 7.5, 50 mM NaCl, with freshly added 1 mM TCEP). Protein elution off the column was induced through a salt gradient between Anion A buffer and Anion B buffer (50 mM Tris pH 7.5, 1 M NaCl, with freshly added 1 mM TCEP). The eluted protein was cleaved at the His6-TEV tag by addition of 0.10 mg/mL of His6-tagged TEV protease at 4 °C overnight. This cleavage cocktail was flowed through a 2 mL Ni-NTA gravity column (ThermoFisher) to separate the cleaved protein from uncleaved protein and TEV protease. Finally, the cleaved protein was purified by size-exclusion chromatography on a Superdex 200 10/300 gel filtration column (Cytiva) equilibrated with SEC buffer (20 mM HEPES pH 7.5, 150 mM NaCl, and 10% glycerol). Pure fractions were pooled and concentrated, and flash frozen in liquid N2 for long-term storage at −80 °C.

### Basal activity measurements

Initial rate measurements for the SHP2-catalyzed dephosphorylation of 6,8-difluoro-4-methylumbelliferyl phosphate (DiFMUP) were conducted at 37 °C in DiFMUP buffer (60 mM HEPES pH 7.2, 150 mM NaCl, 1mM EDTA, 0.05% Tween-20). Reactions of 50 µL were set up in a black polystyrene flat bottom half area 96-well plate. A substrate concentration series of 31.25 µM, 62.5 µM, 125 µM, 250 µM, 500 µM, 1000 µM, 2000 µM and 4000 µM was used to determine $k_{cat}$ and $K_M$. Reactions were started by addition of 2.5 nM of SHP2 wild-type, E139D, and R4A; 1 nM of SHP2 E76K, E76K+E139D, E76K+E139A, and R4A+E139D; and 5nM of all the R5A associated mutants and the rest of the E139A associated mutants. Emitted fluorescence at 455 nm was recorded every 25 seconds in a span of 20 minutes using a BioTek Synergy Neo2 multi-mode reader.

The linear part of the reaction progress curve was determined by visual inspection and fit to a line. Slopes were converted from fluorescence units as a function of time to product formation as a function of time using standard curves measured with the reaction products (6,8-difluoro-7-hydroxy-4-

methylcoumarin). Finally, these rates were corrected for enzyme concentration by dividing the values the concentration of enzyme used in the experiment to yield $V_0$ / [enzyme] in units of ($s^{-1}$). These corrected rates were plotted as a function of substrate concentration and analyzed two different ways. First, we attempted to directly fit the rate versus [substrate] data to the Michaelis-Menten equation via non-linear regression analysis. For most SHP2 variants, this yielded suitable fitted values for $k_{cat}$ and $K_M$ (**Supplementary Table 1**). However, for slower mutants, which generally had weaker $K_M$ values, the fits were poor. As an alternative, we did a double-reciprocal transformation of the data and fit it to a line, to do a Lineweaver-Burk analysis. Here, the individual $k_{cat}$ and $K_M$ often showed variance across replicates, but catalytic efficiency ($k_{cat}/K_M$), derived from the inverse of the slope was reliable. Critically, catalytic efficiency values determined by direct fitting of the Michaelis-Menten equation and those determined by Lineweaver-Burke analysis were highly similar, where both analyses where possible (**Supplementary Table 1**). We opted to use the $k_{cat}/K_M$ values from Lineweaver-Burke analysis throughout the paper for consistency. For all mutants, experiments were generally repeated at least three times, and replicates were individually fit using the methods described above. The average and standard deviation of the fitted catalytic parameters across all replicates were calculated, and these values are reported in **Supplementary Table 1**.

### Differential scanning fluorimetry

Purified SHP2 protein stocks were thawed and diluted in DSF buffer (20 mM HEPES pH 7.5, 50 mM NaCl, 0.4% DMSO). 19 µL of buffer was added to a MicroAmp Fast Optical 96-well Reaction plate (Applied Biosystems, # 4346906). 1 µL of 500x SYPRO Orange Protein Gel Stain (Thermo Fisher, catalog no. S-6650) was added to a final protein concentration of 10 µM and 25x SYPRO Orange. Melting curves were performed in an Applied Biosystems Step-One Plus RT-PCR thermocycler. Temperature measurements started at 15 ºC, and temperature was raised by 0.5 ºC every minute with continuous measurements of fluorescence (excitation: 472 nm; emission: 570 nm). Raw fluorescence values along with corresponding temperatures were analyzed using DSFworld (Wu *et al*, 2024), and $T_m$ values were calculated using dRFU. Average $T_m$ values are reported in **Supplementary Table 2**.

### Cell culture

Cells were cultured in a 37 °C tissue culture incubator with 5% CO2. Cells were discarded by passage 25 and tested for mycoplasma every 3 months. HEK 293 cells were grown in Dulbecco's Modified Eagle Medium (DMEM) with 10% Fetal Bovine Serum (FBS).

### Co-immunopurification

$10^6$ HEK 293 cells were seeded in a 6 cm plate. The next day, cells were transfected using 3 µg of SHP2 DNA, 3 µg of either Gab1,Gab2, or PD-1, 1 µg c-Src and 21 µg PEI in 700 µL DMEM. The transfection medium was refreshed after 16 hours and replaced with DMEM with 10% FBS. After 48 hours, cells were harvested by scraping in PBS. Cells were washed 3 times in 1 mL PBS, and lysed in 100 µL lysis buffer (20 mM Tris-HCl, pH 8.0, 137 mM NaCl, 2 mM EDTA, 10% glycerol, and 0.5% NP-40 + protease inhibitors + phosphatase inhibitors) for 30 minutes on ice. Cells were spun at 17.7 rpm for 15 minutes at 4 °C. Supernatant was transferred to a clean Eppendorf tube and stored at -20 °C.

Protein concentration was determined using a bicinchoninic acid (BCA) assay and absorbance was measured at 562 nm using a BioTek Synergy Neo2 multi-mode reader. 800 µg of protein in a total volume of 380 µL was incubated overnight with 30 µL of magnetic Myc-beads while rotating at 4 °C overnight. The beads were washed 3 times on a magnetic rack using 1 mL lysis buffer. Then, 65 uL1x Laemmli buffer was added and beads were boiled at 100 ºC for 5-8 minutes. For whole cell lysates, 15 µg protein was loaded onto a gel. For IP samples, 15 µL of boiled supernatant was used. Gel was transferred onto a nitrocellulose membrane using TransBlot Turbo (BioRad) and the membrane was blocked using 5% bovine serum albumin (BSA) in Tris-buffered saline (TBS) for 1 hour at room temperature. Membranes were rinsed with TBS with 0.1% Tween-20 (TBS-T) and incubated with primary antibodies in TBST + 5% BSA overnight at 4ºC (Vinculin, CST, #13901S,1:1000; Myc, Invitrogen, #R95025, 1:5000; FLAG, CST, #8146S, 1:1000; pTyr, CST, #8954S, 1:2000). SHP2 was detected using an α-Myc antibody, whereas co-immunoprecipitation of the interacting protein was detected using an α-FLAG antibody. Membranes were washed and incubated with secondary antibodies (IRDye 680 and 800; LiCor #926-68071 and #926-32210, respectively). Blots were imaged on a LiCor Odyssey. Band intensities were quantified using Image Studio Lite (Version 5.2) using the Median Background setting. IP intensities were divided by corresponding intensities in total cell lysate.

Calculation of free energy differences and coupling energies in double mutant cycles

Differences of Gibbs free energy changes between catalysis of SHP2 mutants ($\Delta\Delta G^{\ddagger}$ values) were calculated with the following equation (Carter *et al*, 1984):

$$\Delta\Delta G^{\ddagger} = -RTln\frac{(k_{cat}/K_M)_{mut1}}{(k_{cat}/K_M)_{mut2}}$$

In our calculations, R = 0.001987 kcal · $mol^{-1}$ · $K^{-1}$, T=310.15K for all reactions were conducted under 37°C. $\Delta\Delta\Delta G^{\ddagger}$ values were calculated by the subtraction of the $\Delta\Delta G^{\ddagger}$ values on opposite edges of the double mutant cycles.
